# PM2.5 toxin benzo[a]pyrene induces life-limiting inflammation and oxidative stress in the airway by up-regulation of TRPC6 and inactivation of β2AR/CFTR signaling

**DOI:** 10.64898/2026.04.21.719931

**Authors:** Hung Caohuy, Mungunsukh Ognoon, Tinghua Chen, Qingfeng Yang, Thalia Dib, Bette S. Pollard, Naheed Fatima, Thomas Flagg, Dharmendra K. Soni, Roopa Biswas, William Rittase, Oliver J L’Esperance, Sharon Juliano, Harvey B. Pollard

## Abstract

**Background:** Sustained exposures to high atmospheric levels of PM2.5 at population scale are associated with increased risks for pulmonary inflammatory diseases. These are marked by activation of the TRPC6 (Transient Receptor Potential Canonical type 6) calcium channel, increased reactive oxygen species (ROS) and oxidative stress. Long term exposures are associated with reduced life span, and increased incidences of cardiovascular diseases, dementia, Parkinson’s and Alzheimer disease, and increased risk of autism and autism spectrum disorders. It has been proposed that the PM2.5 toxin is benzo[a]pyrene (B[a]P) that is adsorbed to the surface of the PM2.5 particle.. But the mechanism by which B[a]P might drive pulmonary inflammatory diseases, or any other of the indications above, are not known.

**Hypothesis:** B[a]P was recently reported to bind irreversibly and destructively to the β2 Adrenergic Receptor (β2AR) in the lung. *We* have therefore hypothesized that B[a]P is the adsorbed PM2.5 toxin, and that β2AR is the B[a]P receptor responsible for TRPC6 activation in lung epithelial cells.

**Results:** To test this hypothesis, we exposed a polarized organoid model of normal human lung epithelia, polarized lung epithelial 16HBE14o-cells, and tracheobronchial slice cultures from ferret lung to either PM2.5 or B[a]P. We found that both PM2.5 and B[a]P: (i) irreversibly activated of β2AR signaling via G_i_ to PI3K/AKT; (ii) increased NFκB-activated release of proinflammatory cytokines through IKKαβ activation by PI3K/AKT, which was suppressed by the PI3K inhibitor LY 294002 (iii) desensitized and destroyed the activated β2AR receptor by endocytic recycling; (iv) also destroyed β2AR’s signalplex partner CFTR by the same process; (v) activated the CFTR-bound calcium channel protein TRPC6 due to loss of inhibitory CFTR; leading to (vi) increased cytosolic [Ca^2+^] concentration; (vii) increased ROS due to mitochondrial uncoupling; and (viii) increased expression of oxidative stress. Treatment with the TRPC6 inhibitor BI 749327 blocked steps (vi-viii), and preserved CFTR from endocytic loss. Treatment of tracheobronchial slice cultures of ferret lung with either PM2.5 or B[a]P resulted in increased secretion of IL-6, increased expression of TRPC6, and reduced expression of β2AR and CFTR. Finally, we found that exposure of lung organoids to B[a]P significantly reduced expression of the same five microRNAs (miR-126a-3p, miR-30b-5p, miR-103a-3p, miR-26a-5p, and miR-766-3p) previously identified in sera from service members exposed to PM2.5 from burn pit emissions during deployment to Iraq and Afghanistan.

**Conclusion:** PM2.5 and the PM2.5 toxin benzo[a]pyrene (B[a]P) induce inflammation and oxidative stress in the airway by increased expression of TRPC6 and inactivation of β2AR/CFTR signaling. These discoveries mark the first identification of a mechanism by which exposure to PM2.5 or the PM2.5 toxin B[a]P itself can induce inflammation and TRPC6-dependent oxidative stress in lung epithelia.

## Introduction

Exposure to high atmospheric levels of PM2.5 are associated with increased risks for inflammatory lung diseases, including asthma-like illnesses, <u>c</u>hronic <u>o</u>bstructive <u>p</u>ulmonary <u>d</u>isease (COPD), interstitial lung disease (ILD), and pulmonary fibrosis (PF) [1–4]. Long term PM2.5 exposures are also associated with reduced life span [5, 6], and increased incidences of cardiovascular diseases [7–11], dementia [12–15]), and neurological disorders such as Parkinson’s or Alzheimer’s disease [16, 17]. Reduced lifespan due to exposure to PM2.5 has recently been further characterized by measures of Premature Mortality (PM) and Years of Life Lost (YLL) [18]. Compared to ozone (O_3_) or NO_2,_ PM2.5 was found to be the most significant contributor to PM and YLL. In addition, children exposed to high levels of PM2.5 during the first trimester of gestation are associated with increased risk of autism [19–21] and autism spectrum disorder [22, 23]. The culprit, PM2.5, is <u>p</u>articulate <u>m</u>atter with a diameter of <u>2.5</u> microns or less, that is emitted by both high-heat cooking methods and large-scale incomplete combustion of organic and fossil fuels. The inhaled toxic threshold dose of PM2.5 is ≥ 15 ug/m^3^ [24]. In addition, the small size of PM2.5 means that when inhaled, it can penetrate deeply into the lung parenchyma [1]. There, most of the PM2.5 remains and contributes to lung toxicity [3], where it can induce: (i) inflammation by activation of NFκB signaling and cytokine production; (ii) increased cytosolic Ca^2+^ through the calcium channel protein TRPC6; (iii) mitochondrial dysfunction, involving uncoupling of Complex I and III in the electron transport system; (iv) generation of Reactive Oxygen Species (ROS) and mitophagy; and (v) oxidative stress, leading to DNA strand breaks, protein oxidation, membrane disintegration and cell death [6, 25–27].

However, a small fraction can also enter the blood stream through the alveolar-capillary barrier [1, 3], and thereby reach the heart [28], brain [29] and placenta [30]. Consistently, similar proinflammatory and oxidative stress consequences have been reported for the heart [31–33] and for the brain [34, 35]. *Nonetheless, whether in the lung or elsewhere in the body, the mechanisms initiating PM2.5’s inflammatory and oxidative sequelae remain unknown, and the development of candidate therapies has therefore languished*.

The core of the PM2.5 particle itself is amorphous silicon dioxide (SiO_2_), which has been classified by the FDA as GRAS, meaning <u>G</u>enerally <u>R</u>ecognized <u>a</u>s <u>S</u>afe for inclusion in food. and as safe by the European Food Safety Authority (EFSA: Code E551). When eaten, very little amorphous SiO_2_ enters the blood stream, and thus resembles dietary fiber [36]. Conceptually, plants contain up to 10-15% amorphous SIO_2_, which is why incompletely combusted organic matter contains this substance [37]. Experimentally, extraction of PM2.5 by sonication in aqueous buffers releases lipopolysaccharides (LPS) and other inorganic substances, including transition metals and heavy metals [38]. Thereafter, extraction with organic reagents removes toxic polyaromatic hydrocarbons (PAH), leaving behind a benign residual particle [39].

Consequently, investigation has shifted to potentially toxic co-combustion products that are adsorbed to the surface of the PM2.5 core particle [40, 41]. Among these, it has been reported that an important PM2.5 toxin may be Benzo[a]Pyrene (B[a]P), a member of the class of polyaromatic hydrocarbons (PAH) which are emitted during incomplete combustion of organic matter [42]. In earlier studies, B[a]P was calculated to be responsible for up to 67% of the PAH-based toxicity associated with PM2.5 [43]. More recently, B[a]P has been described as “the most toxic combustion product of fossil fuels known to man.” [41, 44–46]. *However, despite extensive evidence linking B[a]P exposure to pulmonary inflammation, the underlying mechanisms by which B[a]P engages intracellular signaling pathways remain obscure,* Given the lack of mechanistic insight, we speculated that B[a]P might exert its pathological influence through a receptor capable of triggering TRPC6 activation. One historical possibility was the aryl hydrocarbon receptor (AhR) [47]. As part of the conversion of B[a]P to the carcinogen benzo[a]pyrene diol epoxide (BDPE), AhR binds B[a]P and transfers it to CYP1A1 or CYP1B1 isoforms of cytochrome P-450 in the liver and elsewhere [48, 49]. The BDPE product can then bind to DNA and promote mutagenesis, leading over the long term to lung and other cancers [50, 51]. But while carcinogenesis at population scale is a long-term effect which may be potentiated by chronic airway inflammation and oxidative stress, the PM2.5-dependent activation of proinflammatory NFκB signaling and oxidative stress in the lung appear to be a consequence of a shorter exposure time. Thus, the two effects may be related but are separated over time. However, it has also been suggested that AhR might also directly drive NFκB-mediated proinflammatory B[a]P toxicity [47]. Nonetheless, the data showing activation of NFκB signaling by a hypothetical B[a]P-AhR complex have been conflicting. For example, reports have shown that NFκB signaling increases, decreases, results in a different action [52], or has no effect, depending on specific types of cells [53–55]. Consequently, although AhR does bind B[a]P, the possibility of coincident crosstalk with an AhR-independent biology has been increasingly anticipated [56–58]. In addition, while some studies have shown increases in NFκB signaling, none have shown that a B[a]P complex with AhR can either increase TRPC6 expression or drive TRPC6-dependent calcium influx. *Therefore, while it was clear that AhR does bind B[a]P, and does orchestrate important steps in carcinogenesis, it was also possible that there might also be another B[a]P interacting protein that was independent of AhR and activated TRPC6*.

As an alternative candidate for the TRPC6-activating B[a]P receptor we have considered the beta-2 adrenergic receptor (β2AR), which was recently reported to bind B[a]P irreversibly to its catecholamine binding site [59]. Because B[a]P cannot dissociate from the binding site, B[a]P initially behaves as a super agonist. But, soon thereafter, a fraction of the B[a]P:β2AR complex begins to physiologically desensitize by undergoing endocytic recycling and destruction in the proteosome. Interestingly, in lung epithelial cells β2AR is bound to CFTR, the Cystic Fibrosis (“CF”) gene product, by means of a PDZ domain-mediated signalplex, which is linked together by the PDZ adaptor protein NHERF1 [60]. Native CFTR forms a dimer in the membrane and thus can bind at least two PDZ-containing protein ligands. For example, one of these could be β2AR, and the other could be TRPC6. When functional CFTR is missing, as for people with CF who have two disease-causing CFTR variants, the lack of functional CFTR results in (i) activation of proinflammatory TNFα/NFκB signaling and elevation of the cytokines IL-6 and IL-8 [61]; (ii) increased expression of the calcium channel protein TRPC6 that had been kept in an inhibited state in a signalplex with CFTR [62]; (iii) elevated cytosolic calcium from the TRPC6 channel [62]; (iv) production of increased ROS from distressed mitochondria; and (vi) a state of increased oxidative stress [63, 64]. Consistently, when CFTR is reduced in the larger population of people who have two functional CFTR genes, TRPC6 expression and TRPC6 channel activity are once again elevated, and cytosolic calcium concentration is chronically elevated [65]. In lung epithelia and other cell types, exposure to PM2.5 induces initial increases in cytosolic [Ca2+] from store operated Orai/STIM channels [66]. But subsequently much greater amounts of [Ca2+] enter the cytosol from outside the cell by activated TRPC6 channels [67–69]. In addition, bronchial epithelial cells respond to classical oxidants like H_2_O_2_ by reduction in CFTR and by robustly increased TRPC6 expression [69]. Thus, in the larger non-CF population, TRPC6 may mediate PM2.5-induced ROS and oxidant stress. Nonetheless, the mechanism by which PM2.5 activates TRPC6 is unknown. PM2.5 exposure has also been shown to elevate TRPC6 channel activity due to phospholipase C-dependent action on PIP2 and generation of DAG, although the initiating mechanism is also not known [70–72]. However, when excess β2AR agonist causes desensitization by removal of β2AR from the plasma membrane by endocytic recycling, CFTR channel activity is also reduced by *ca.* 60% [73]. Under similar circumstances, CFTR surface protein expression was reported to decrease by *ca*. 50% [74]. Thus, β2AR abundance may exert control over CFTR abundance at the cell surface. Consequently. CFTR loss could be involved in TRPC6 elevation in response to activation and loss of β2AR. *Consistently, our initial preliminary data in a differentiated lung organoid model revealed that B[a]P, by irreversibly activating both G_s_ and G_i_ mediated β2AR activities, caused loss of both β2AR and CFTR proteins, loss of cAMP-activated CFTR chloride channel activity, and increased TRPC6 expression*.

Based on these and other preliminary data, *we* have hypothesized that B[a]P is the adsorbed PM2.5 toxin, and that the B[a]P receptor responsible for TRPC6 activation in lung epithelial cells is β2AR. Thus, it is possible that the B[a]P-induced activation and then loss of both β2AR and CFTR, and consequent elevation of TRPC6, might explain the pulmonary pathology caused by PM2.5 exposure. To test this hypothesis we proposed to determine whether parallel studies of B[a]P and PM2.5 resulted in equivalent effects on lung pulmonary epithelial cells by: (i) initially activating the β2AR signaling pathway through the G_i_ protein to activate the proinflammatory [PI3K/AKT/IKKαβ/NFκB] pathway; (ii) thereby activating expression of cytokines IL-8 and IL-6, which could be blocked with the PI3K blocker LY-294002; (iii) reducing β2AR and CFTR protein expression over a 24 hour period; (iv) increasing calcium influx through TRPC6 channels following dissociation of TRPC6 from CFTR and by PLC-dependent production of DAG; and (v) and producing Reactive Oxygen Species (ROS) and oxidative stress which could be blocked by the TRPC6 inhibitor BI 749327. Based on the results shown below, we have concluded that the hypothesis is not disproved. These discoveries mark the first identification of a mechanism by which exposure to PM2.5 or the PM2.5 toxin B[a]P itself can induce inflammation and TRPC6-dependent oxidative stress in lung epithelia.

## Results

### B[a]P activates proinflammatory TNFα/NFκB signaling in airway epithelia

To test if incubation of differentiated airway epithelial organoid cultures with B[a]P activated TNFα/NFκB signaling, we incubated BCi.NS1.1 (dBCi) cells differentiated in an air-liquid-interface (ALI) culture with different concentrations of B[a]P for 6 hours apically and extended incubation for additional 18 hours. As previously described [75], we then measured expression levels of a comprehensive set of proteins in the TNFα/NFκB signaling pathway which we expected to be elevated. **Fig 1a** showed that the concentrations of B[a]P chosen for the analysis, 0.1, 1.0 and 10 μM were not toxic to the cells. **Fig 1b** also showed that as the concentrations of B[a]P rose, expressions of IL6, IL8 and TNFα in the liquid subphase also rose. At the lowest 0.1 μM dose of B[a]P, only increases in IL-8 and IL-6 secretion were significant, while at 1 and 10 µM B[a]P all IL-6, IL-8 and TNFα were significantly and proportionately elevated. Finally, **Fig 1c** showed that as B[a]P concentration rose, protein expression also increased for TNFR1, TRADD, p-IKKBαβ, p-IκBα, p-NFκBp65 (serine 536), and p-NFκBp65 (serine 276). By contrast, non-phosphorylated substrate proteins, IKKαβ, IκBα and NFκBp65 were not significantly changed. These data thus showed that B[a]P was activating the classical activation pathway for NFκB. The B[a]P-dependent elevation of TRADD further established that B[a]P was dose-dependently elevating signaling specifically in the TNFα/NFκB pathway.

**Figure 1.**
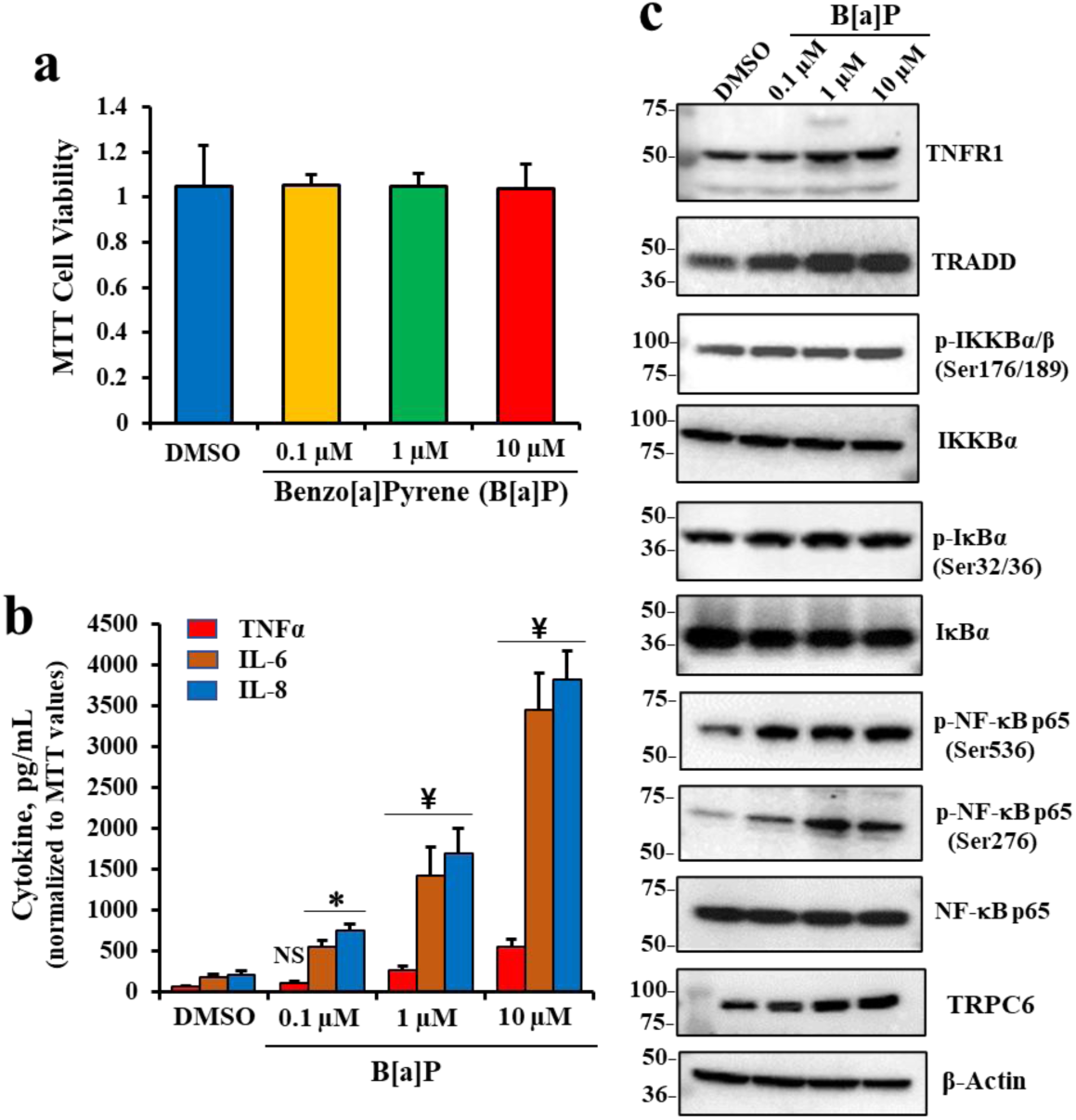
Effects of B[a]P on cell viability and cytokine secretion, and expression of pro-inflammatory signaling proteins in differentiated BCi.NS1.1 (dBCi) cells. (**a**) Cell viability was assessed with the MTT assay and data shown are means ± SD (n = 3). (**b**) TNFα, IL-6 and IL-8 expression in response to B[a]P exposure. (**c**) Representative Western blot images showing protein expression. All data were obtained from three independent experiments. Statistical *p* values were determined by a one-way ANOVA, followed by a Tukey post-hoc test comparing the B[a]P treatments to the DMSO control; * *p* < 0.05; ¥, *p* < 0.01; NS, no significance.

### B[a]P reduces CFTR chloride channel activity and β2AR and CFTR protein expression

Using the Ussing Chamber method to measure cAMP-activated CFTR channel function, **Fig 2a** and **Fig 2b** showed that CFTR channel function, represented by short circuit currents (ΔI_sc_), were systematically and significantly reduced by increases in B[a]P concentrations. Western blots and densitometric analyses shown in **Fig 2c** and **Fig 2d**, respectively, showed that protein expression of both β2AR and CFTR were also significantly reduced by step-wise increases in B[a]P concentration. Finally, **Fig 2e** showed that there was a significant positive correlation between changes in β2AR and CFTR with a Pearson Correlation Coefficient (r) of 0.9948 (p= 0.005). Similar B{a]P effects were observed in polarized lung 16HBE14o-cells (see **Supplemental Fig S1**). *Thus, over 24 hours of incubation, irreversible binding of B[a]P to β2AR results in loss of CFTR channel function and coordinated loss of both β2AR and CFTR proteins*.

**Figure 2.**
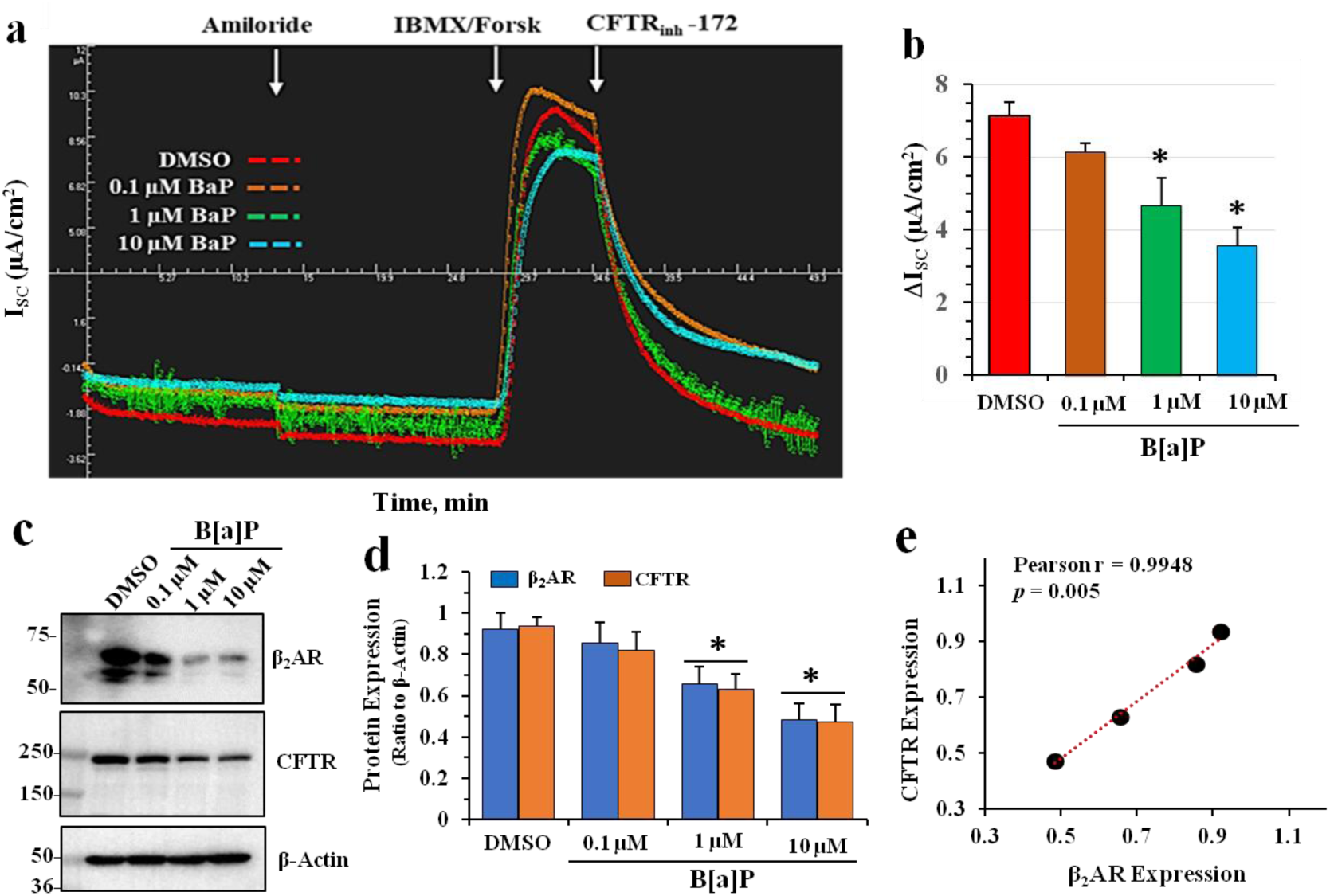
Effects of B[a]P on β2AR and CFTR in differentiated BCi.NS1.1 (dBCi) cells. **(a & b)** Representative *I*_sc_ tracings of CFTR-dependent short-circuit currents (*I*_sc_) were measured in Ussing chambers. **(c)** Representative Western blot images of total cellular CFTR and β2AR expression levels in treated dBCi cells after Ussing chamber analyses in (**a**). **(d)** Densitometric analyses showing B[a]P-induced reduction of both β2AR and CFTR expression in a dose-dependent manner. **(e)** The Pearson correlation coefficient (r) showing a significant positive relationship between B[a]P-inhibited β2AR and CFTR expression. All data were obtained from three independent experiments. Statistical *p* values were determined with a one-way ANOVA, followed by the Tukey post-hoc test comparing the B[a]P treatments to the DMSO control; * *p* < 0.01.

### PM2.5 and B[a]P increase cytokine expression and proinflammatory signaling

To test whether the parent PM2.5 exerted similar effects to B[a]P on lung epithelial cells, we subjected polarized 16HBE14o-cells to different concentrations of authentic PM2.5. The PM2.5 we utilized was ERM-CZ110 from Sigma-Aldrich (see Methods for additional details). We also included 10 μM B[a]P as a positive control. **Fig 3a** showed that the concentrations of PM2.5 chosen for the analysis, 12.5, 25, 50 and 100 ng/ml, were not toxic to the cells. **Fig 3b** showed that as the concentrations of PM2.5 rose, expressions of IL6 and IL8 in the liquid subphase also rose. Finally, **Fig 3c** showed that as PM2.5 concentration rose, expression of both CFTR and β2AR progressively declined, as previously shown for pure B[a]P (**Fig 2c** & **2d**) and in the 10 μM B[a]P positive control. PM2.5 was also found to significantly reduce cAMP-dependent CFTR chloride channel activity (see **Supplemental Fig S2**). **Fig 3c** also showed, as anticipated from initial activation of β2AR by the irreversible binding of B[a]P, that both the B[a]P positive control and PM2.5 increased p-AKT (ser 473) expression. In addition, proinflammatory p-NFκBp65 (serine 536) was elevated by both 100 ng/ml PM2.5 and the B[a]P positive control. Finally, we noted a progressive PM2.5-dependent increase in the calcium channel protein TRPC6 and for the B[a]P positive control. Normally, CFTR binds to and constitutively inhibits the TRPC6 calcium channel. However, in the absence of CFTR (*vide supra*) TRPC6 is activated and further expressed in the apical plasma membrane, thereby contributing to oxidative stress [62]. *These data thus show that the effects of PM2.5 and the pure B[a]P are shared for (i) loss of β2AR and CFTR expression; (ii) activation of AKT; (iii) activation of proinflammatory NFκB signaling; and (iv) increased expression of TRPC6*.

**Figure 3.**
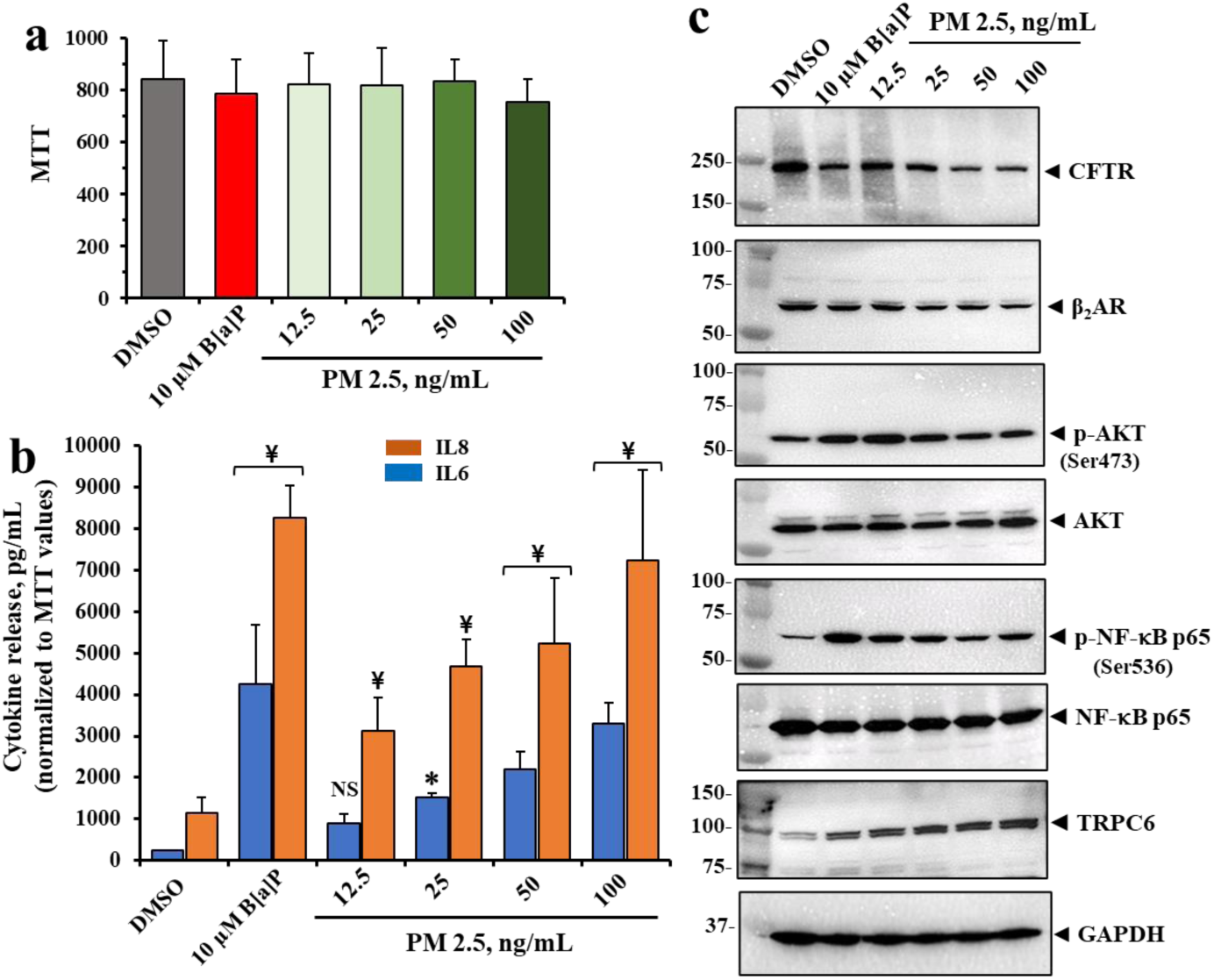
Effects of PM 2.5 on cell viability and cytokine secretion, and expression of pro-inflammatory signaling proteins in polarized 16HBE14o-cells. Cells were treated with various concentrations of PM2.5 for 24 hr. (**a**) Cell viability was assessed with MTT assay. (**b**) IL-6 and IL-8 from the basolateral sides. (**c**) Representative Western blot images showing CFTR, β2AR, phospho-AKT (Ser473), AKT, phospho-NF-κB p65 (Ser536), NF-κB p65, and TRPC6 expression. All data are presented as the mean ± SD (n = 4). Statistical *p* values were determined with a one-way ANOVA, followed by the Tukey post-hoc test comparing the B[a]P or PM 2.5 treatments to the DMSO control; *, *p* < 0.05; ¥, *p* < 0.01; NS, no significance.

### B[a]P induces endosomal loss of β2AR and CFTR

Plasma membrane proteins like β2AR and CFTR are continuously recycled by uptake into endosomes where the damaged proteins are retained and sent for disposal in lysosomes or proteosomes, and the fully functional proteins are returned to the plasma membrane [75, 76]. To test for whether the formation of irreversible B[a]P/β2AR complexes lead to endosomal loss of both β2AR and CFTR, we labeled both membrane-localized β2AR and CFTR with impermeant biotin, and followed the endosomal processing of both proteins. First, cells were treated for 6 hours with different concentrations of B[a]P and then were labeled at 4°C with impermeant Sulfo-NHS-SS-biotin. **Fig 4a** showed that following B[a]P treatment, the detection of surface-labeled biotinylated CFTR and β2AR progressively declined. To test whether B[a]P treatment resulted in both CFTR and β2AR being retained in intracellular endosomes, we pre-labeled cell surface β2AR and CFTR with impermeant Sulfo-NHS-SS-biotin at 4°C for one hour, and then incubated the labelled cells with different concentrations of B[a]P for 6 hours. Thereafter the remaining biotin on surface-localized CFTR or β2AR were stripped with glutathione (GSH), and the “damaged” endosomal biotinylated β2AR and CFTR that was protected within the cell were then identified by Western blot analysis. **Fig 4b** showed that as the concentration of B[a]P increased, progressively more biotinylated β2AR was retained in intracellular endosomes. **Fig 4c** showed the exact same relationship was found for CFTR. These data suggest the irreversible binding of B[a]P to β2AR may have initially resulted in activation of β2AR signaling [59]. But soon thereafter the classical desensitization process begins, extending for hours or days with sustained exposure [77]. Desensitization results in removal of β2AR from the plasma membrane, retention in endosomes, and eventual proteosomal destruction. We find here that CFTR is also removed from the plasma membrane by the same or parallel process, specifically involving proteosomal destruction (**Fig 4d**)*. Thus, the formation of the irreversible B[a]P/β2AR complex leads to activation of β2AR signaling, followed by endosomal loss of both β2AR and CFTR*.

**Figure 4.**
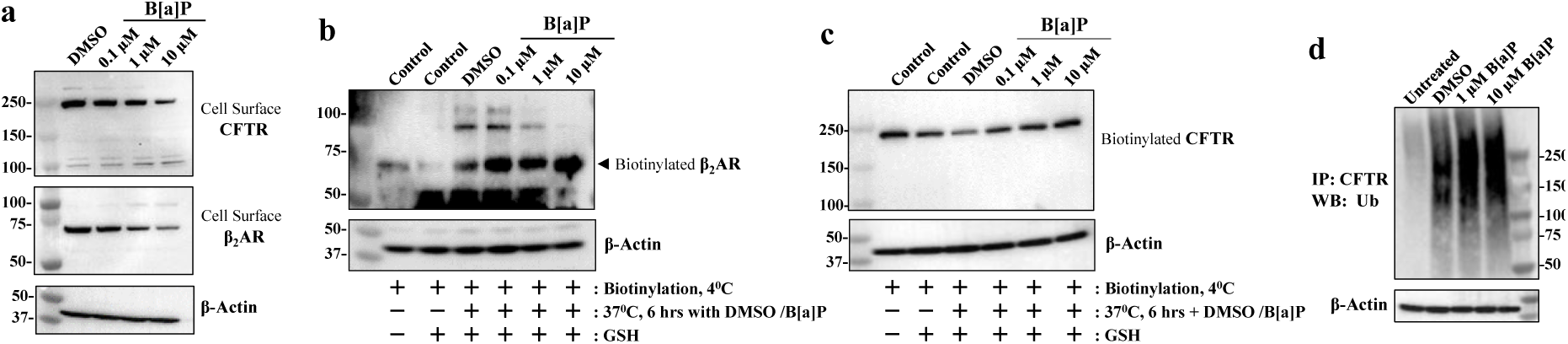
B[a]P induces loss of cell surface β2AR and CFTR expression by inhibiting endosomal recycling in differentiated BCi.NS1.1 (d-BCi) epithelia. **(a)** Representative Western blot images of cell surface CFTR and β2AR expression in cells treated with DMSO or various concentrations of B[a]P. **(b & c)** Recycling assay of endocytosis CFTR and β2AR were analyzed by Western blot. Representative Western blot images of biotinylated CFTR and β2AR expression are shown. Cell samples treated without or with GSH (*lanes 1 and 2*) were used as positive controls for biotinylation and GSH stripping processes, respectively. **(d)** A representative Western blot image showing ubiquitination of CFTR immunoprecipitates from cell lysates treated with different concentrations of B[a]P in the presence of 20 µM MG 132. All Western blot analyses represent the results of three independent experiments.

### PM2.5 and B[a]P induce cytokine secretion by activating the PI3K/AKT/IKKαβ/NFκB pathway

As shown in **Figure 2a**, β2AR can couple canonically with G_S_ proteins to activate adenyl cyclase, which then can activate protein kinase A to phosphorylate and activate the CFTR chloride channel [78]. However, β2AR can also couple non-canonically with G_i_ proteins by a pathway, modulated by sirtuins 1 and 3, that lead to activation of the [PI3K (phosphoinositide 3 kinase)/AKT (protein kinase B)/IKKαβ/NFkBp65] pathway [79]. Both PM2.5 (**Fig 3c**) and pure B[a]P (**Fig 1c**), can activate p-AKT signaling and p-NFkBp65 (Ser 536). Thus, B[a]P-activated β2AR may activate PI3K through G_i_, and thus activate the remainder of the [PI3K/AKT/IKKαβ/NFκBp65] pathway. To test this hypothesis, we asked whether the classical PI3K blocking drug LY 294002, but not its inactive analogue LY 303511, could block B[a]P-activated cytokine expression by lung epithelial cells. **Fig 5a** showed that LY 294002 blocked B[a]P-activated secretion of IL-8 and IL-6, However, the inactive form LY 303511 was inactive. These data suggested that B[a]P could activate NFκB-dependent secretion of the cytokines IL-8 and IL-6 by activation of the [β2AR/G_i_/PI3K/AKT/IKKαβ/NFκB] pathway. Equivalent experiments with PM2.5 treated cells, incubated with LY 294002 or the inactive analogue LY 303511 were also tested with similar statistically significant results (**Supplemental Fig S3**). *Thus, B[a]P] can activate proinflammatory p-NFkBp65 signaling in two ways: (i) through irreversibly activating β2AR-dependent PI3K signaling, and (ii) by inactivating CFTR, thereby relieving inhibition of TRADD. Both pathways converge on IKKαβ to commonly phosphorylate IκBα and free p-NFkBp65 to leave the cytosol and seek out kB binding sites on cytokine promoters within the nucleus*.

**Figure 5:**
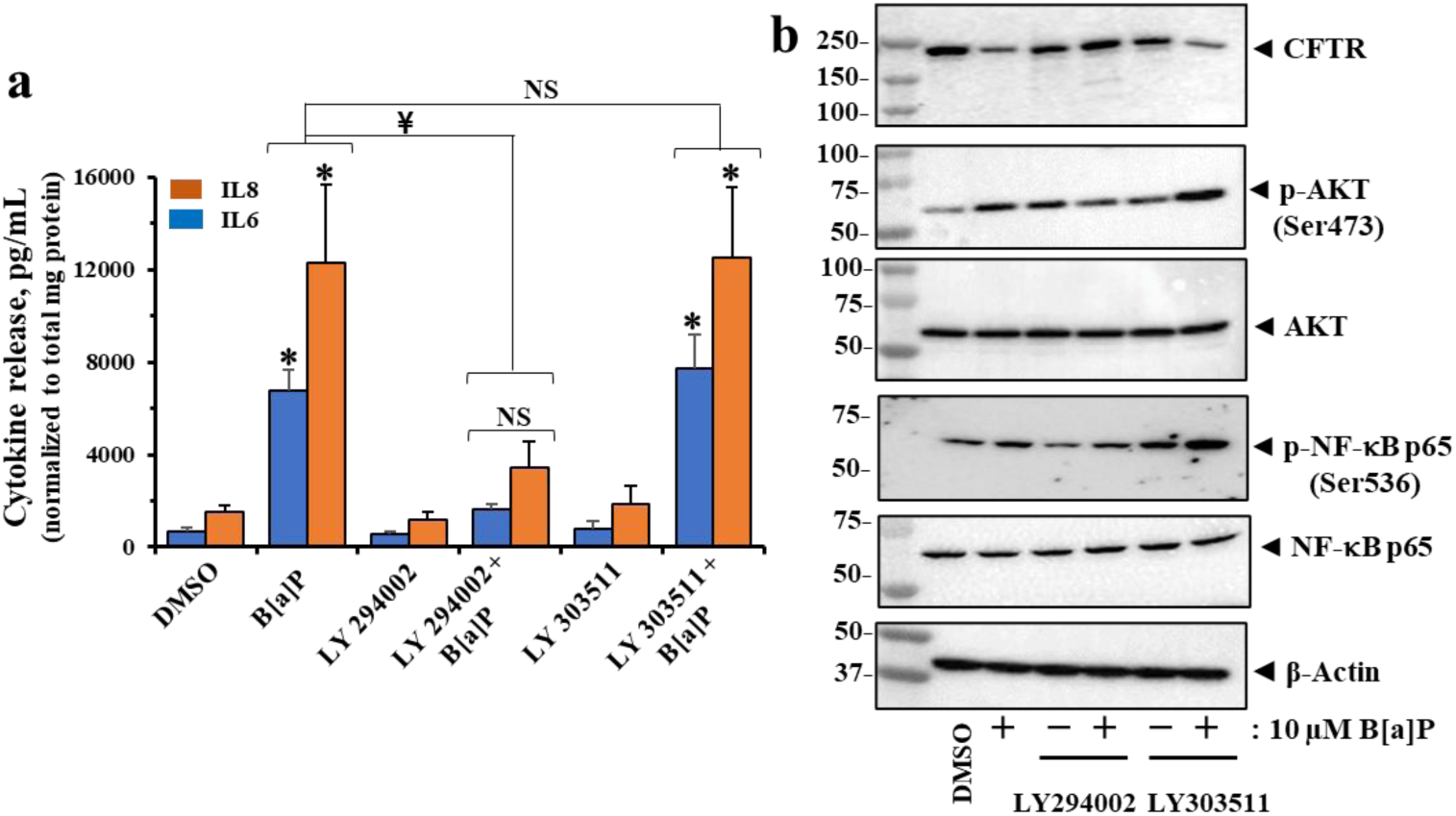
B[a]P induces cytokine secretion via the PI3K signaling pathway in polarized 16HBE14o-cells. (**a**) IL-6 and IL-8 from the basolateral sides of the ALI were determined using the DuoSet® ELISA kits. Data are presented as the mean ± SD (n = 4). Statistical *p*-values were determined with a one-way ANOVA, followed by the Tukey post-hoc test comparing the all treatments to the DMSO control; ¥, *p* < 0.05; *, *p* < 0.01. (**b**) Representative Western blot images showing CFTR, phospho-AKT (Ser473), AKT, phospho-NF-κB p65 (Ser536), and NF-κB p65 expression in treated polarized 16HBE14o-cells in (**a**). All data were obtained from three independent experiments.

### PM2.5 and B[a]P activate calcium influx through TRPC6 channels

As shown in **Fig3c**, we also determined that PM2.5 exposure to lung epithelial cells induced elevation of TRPC6 expression. Therefore, to test whether exposure to B[a]P activated calcium influx through TRPC6 channels, we exposed cells to different concentrations of B[a]P for 24 hours, and then treated cells with calcium Fluo-4 AM. Changes in intracellular calcium concentration sere then measured following exposure to 1 mM extracellular calcium concentration. **Fig 6a** & **6b** showed that after 24 hours of treatment a significant increase in intracellular calcium could be measured with 1 and 10 μM B[a]P. **Fig 6c** & **6d** showed that B[a]P significantly and dose-dependently reduced CFTR and β2AR, and elevated TRPC6 expression. Equivalent results to those in **Fig 6c** & **6d** were also found when performed with PM 2.5 instead of B[a[P (see **Supplemental Fig S4**). **Fig 6e** showed that when comparing loss of β2AR to increase in TRPC6, the Pearson’s r was -0.9875 (*p* = 0.01). Finally, to test directly whether calcium intake occurred through TRPC6 channels, we repeated the calcium measurement experiments in the presence of the specific TRPC6 inhibitor BI 749327. **Fig 6f** & **6g** showed that BI 749327 significantly inhibited B[a]P-dependent calcium influx. Then, to test whether BI 749327 affected stability of CFTR following exposure to B[a]P, Western blot analysis in **Fig 6h**, prepared from cells in **Fig 6f** & **6g,** revealed that BI 749327 significantly protected CFTR from loss due to exposure to B[a]P. Thus, stabilization of CFTR by the TRPC6 inhibitor BI749327 prevented B[a]P and PM2.5 from activating the TRPC6 channel. *Altogether these data suggested that activation of TRPC6 channels by PM2.5 was due to the B[a]P adsorbed to the particle surface, and that the TRPC6 inhibitor BI 749327 was able to preserve the integrity of the complex between CFTR and TRPC6 from destruction by B[a]P*.

**Figure 6.:**
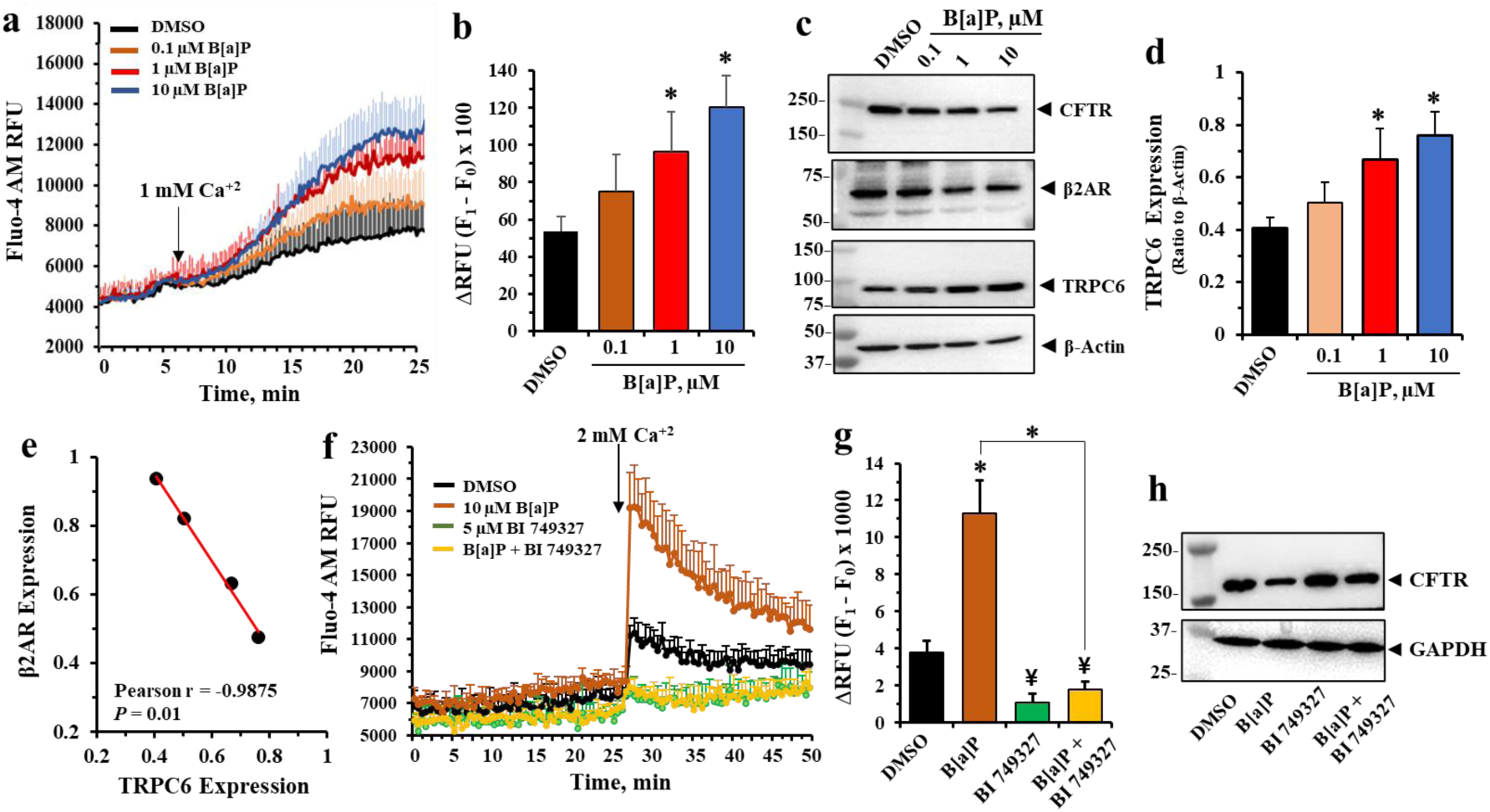
B[a]P induces calcium influx which is blocked by TRPC6 inhibitor BI 749327 in 16HBE14o-cells. (**a**) Fluo-4-AM fluorescence intensity traces obtained 24 hours after treatment with B[a]P and stimulated with assay buffer containing 1 mM Ca^+2^. (**b**) Corresponding histograms summarize the mean changes in RFU (F_1_ – F_0_) after stimulation by 1 mM Ca^+2^. (**c** & **d**) Representative Western blot images of β2AR and TRPC6 expression levels in treated cell in (**a**) and respective densitometric analyses. (**e**) The Pearson correlation coefficient (r) showing a significant negative relationship between β2AR and TRPC6 expression caused by B[a]P treatment. (**f**) Fluo-4AM fluorescence intensity traces in 16HBE14o-cells were obtained before and after addition of 2 mM Ca^+2^ in the presence or absence of 5µM BI 749327. (**g**) Corresponding histograms summarize the mean changes in RFU (F_1_ – F_0_) after stimulation by 2 mM Ca^+2^. (**h**) Representative western blotting analyses of CFTR expression in corresponding treated cells. All data represent the mean of three independent experiments. Statistical *p* values were determined with a one-way ANOVA, followed by the Tukey post-hoc test comparing the B[a]P ± BI 749327 treatments to the DMSO control; ¥, p<0.05, * *p* < 0.01.

### PM2.5 and B[a]P induce production of Reactive Oxygen Species (ROS) and Oxidative stress

By inducing Ca^2+^ influx through the TRPC6 calcium channel due to loss of CFTR control, and by availability of DAG from activation of the β2AR/PI3K/PLC pathway, we expected that increased ROS and oxidative stress might follow. To test this expectation, we treated cells with two incrementing doses of PM2.5, or one dose of B[a]P for 24 hr, and then assayed them for ROS, cytokine release and CFTR expression. We also included 100 µM H_2_O_2_ as a positive control for oxidative stress. **Fig 7a** showed that 50 and 100 ng/ml of PM2.5, or 10 μM B[a]P, or 100 μM H_2_O_2_, significantly increased generation of ROS. **Fig 7b** showed that IL-6 and IL-8 were significantly elevated in the same manner as increases in ROS. These results may therefore implicate the Nrf2/Keap1 redox signaling pathway in CFTR repression under oxidative conditions [80]. **Fig 7c** showed that CFTR expression was significantly reduced by exposures to both PM2.5 and B[a]P. The H_2_O_2_ positive control also significantly lowered CFTR expression, as expected from activation of the distal antioxidant element (ARE) in the CFTR locus [80].

**Figure 7.**
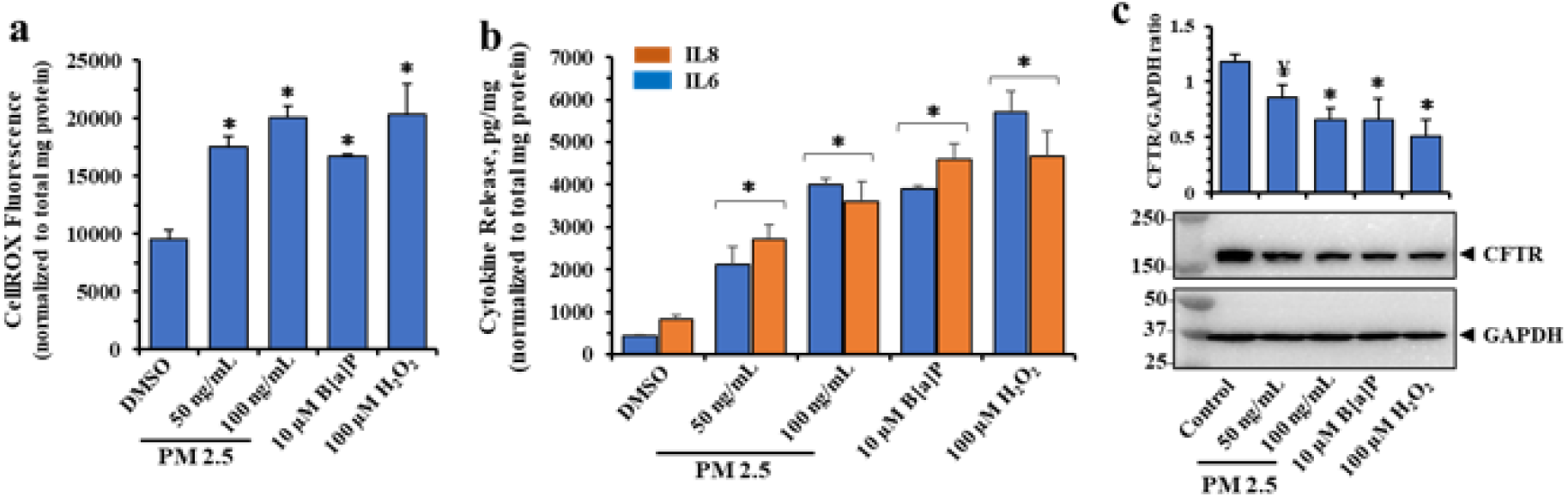
PM 2.5, B[a]P and hydrogen peroxide (H_2_O_2_) induce Reactive Oxygen Species (ROS) production in polarized 16HBE14o-cells. Both apical and basolateral sides of polarized 16HBE14o cells were treated with DMSO, PM 2.5, B[a]P, or H_2_O_2_ (as a positive control for oxidation stress). After 6 hr incubation, treated media from the apical sides only were removed and replaced with HBSS buffer containing 10 µM CellROX Green, followed by further incubation for 18 hrs. (**a**) ROS production significantly elevated by all three analytes. (**b**) Measurement of IL6 and IL8 release in response to all three analytes (**c**) Representative western blot images of treated cell lysates showing significant losses of CFTR under these conditions. Data are presented as the mean ± SD (n = 4). Statistical *p* values were determined with a one-way ANOVA, followed by the Tukey post-hoc test comparing PM2.5, B[a]P, and H_2_O_2_ treatments to the DMSO control; ¥, *p* <0.05; *, *p* < 0.01.

### *In vivo* human exposure to PM2.5 and *in vitro* exposure to B[a]P significantly suppress the same five microRNAs

We previously reported that sera from service members exposed to PM2.5 from open air burn pits exhibited significant and specific reductions in five microRNAs [81]. These five reduced serum microRNAs were: miR 126a-3p, miR-30b-5p, miR103a-3p, miR26a-5p and miR-766-3p, which were identified from among 754 tested microRNAs by a Taqman-based assay. The samples were from pre- *vs*- post- deployment blood draws, and the controls in these studies were service members who had not been deployed. Levels of PM2.5 as high as 615 μg/m^3^ in burn pit emissions were documented as key exposure metrics in this exposed population [82, 83]. Sera from exposed service members were also reported to contain high levels of the B[a]P metabolite BDPE, thus indicating that they had been exposed to B[a]P in the PM2.5 emitted from the burn pits [84]. Consistently, high levels of B[a]P were also measured in serum that had been collected from the service members just after returning from deployment [85]. Therefore, to test whether these *in vivo* results could be replicated *in vitro* by exposure of the lung epithelial organoid to B[a]P, we incubated the apical and basolateral sides of differentiated BCi.NS1.1 organoids with different concentrations of B[a]P and used Taq-man Assays to measure microRNAs. **Fig 8** showed that all five microRNAs were significantly and dose-dependently reduced by exposure to B[a]P. *These data thus suggest that B[a]P effects on microRNA expression, in vitro, can parallel in vivo PM2.5 effects on serum microRNAs in vivo*.

**Figure 8.**
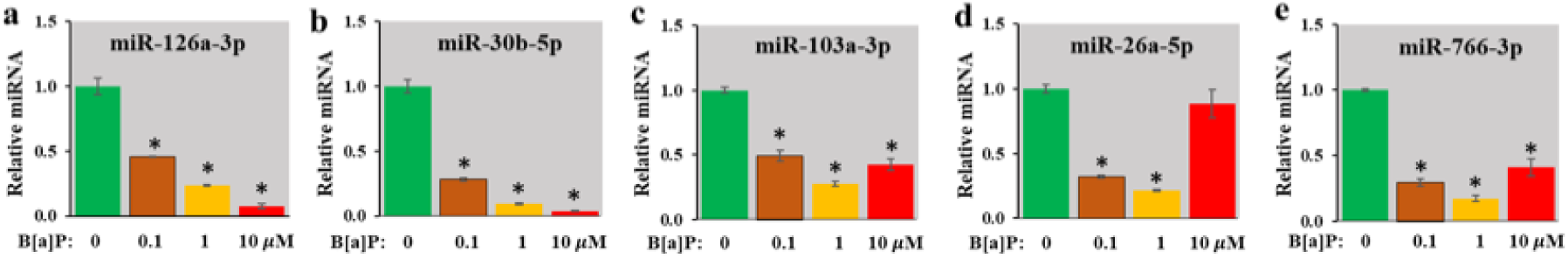
Alterations in miR expressions in response to B[a]P. Both apical and basolateral sides of differentiated dBCi cells were incubated with different concentrations of B[a]P for 24 hours; treated media from the apical sides were removed after 6-hr incubation, and cell treatment continued further under ALI conditions. The cells were lysed and the isolated RNA was analyzed for the expressions of the indicated miRs using miR-specific Taqman Assays (Life Technologies). A dose dependent reduction in the miR levels was observed compared to control cells (green bar). U6 was used as endogenous control. *, *p* < 0.05.

### B[a]P and PM2.5 reduce β2AR and CFTR, and elevate IL-6, and TRPC6 expression in *ex-vivo* Ferret lung tracheo-bronchial slice cultures

To test for the generality of PM2.5 and B[a]P effects on reduced β2AR and CFTR, and for elevation of IL-6 and TRPC6, we turned to the ferret lung. Ferrets are among very best of the small animal models of CF because of their ability to closely phenocopy human CF disease [86, 87]. To test whether the effects of PM2.5 and the PM2.5 toxin B[a]P could be replicated in ferret lung, we prepared tracheo-bronchial slice cultures from ferret lungs and tested whether ex-vivo exposure to PM2.5 or pure B[a]P for 24 hours would not only reduce expression of either β2AR or CFTR, but also activate secretion of IL-6 and increase expression of TRPC6. **Fig 9a** shows results of dissections from two lungs in which the tracheal connections to bronchial bifurcations are identified by yellow arrows. The slices were then cultured as described in the Methods for 24 hours. **Fig 9b** showed that exposure to either 10 μM B[a]P or 100 ng/ml PM2.5 resulted in significant and similar elevations in IL-6. **Fig 9c** showed that both β2AR and CFTR were reduced, while TRPC6 expression was elevated. *These ex-vivo ferret data thus replicated the parallel PM2.5 and B[a]P results shown earlier in differentiated human lung epithelial organoids and polarized lung epithelial cells*.

**Figure 9:**
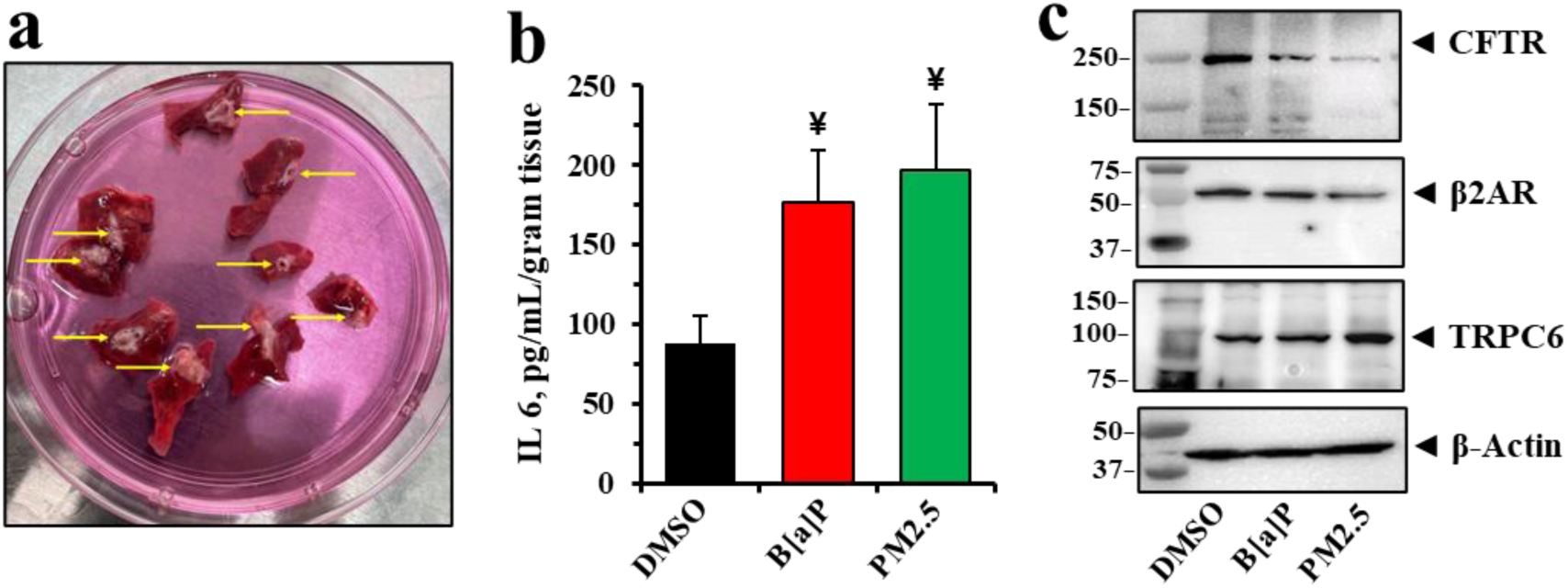
Effects of B[a]P and PM 2.5 in ferret lung tissue *ex vivo*. (**a**) The lung parenchyma from two ferrets were sliced into several sections as shown. *Arrows* indicate the tracheobronchial tubes. All tissue sections were incubating with DMSO (control), 10 µM B[a]P, or 100 ng/mL PM 2.5 in Pneumacult-EX Plus Basal media for 24 hrs. (**b**) IL 6 secretion was determined using the ferret ELISA kit. Data represent the mean of 6 independent sections. Statistical *p* values were determined with a one-way ANOVA, followed by the Tukey post-hoc test comparing the B[a]P or PM 2.5 treatments to the DMSO control; ¥, *p* < 0.05 (n = 6). (**c**) Representative western blotting images show reduction of CFTR, and β2AR and elevation of TRPC6 expression in explants under these treatments.

## Discussion

The data reported here are consistent with the hypothesis that the principal pulmonary PM2.5 toxin is B[a]P, and that β2AR is the B[a]P receptor in lung epithelial cells which is responsible for (i) proinflammatory signaling; (ii) loss of CFTR to activate the TRPC6 calcium channel, and thus for (iii) increases in ROS and oxidative stress. In support of this concept, and as summarized in **Fig 10**, we have shown that both PM2.5 and B[a]P initially activate the β2AR signaling pathway through the G_i_ protein to drive expression of the [PI3K/AKT/IKKαβ/NFκB] pathway. This results in increased expression of IL-6 and IL-8. We validated the increased pathway activity by blocking it with the PI3K blocker LY 294002. Subsequently, apical membrane β2AR and CFTR protein expression declined by endosomal recycling. The loss of CFTR then activated proinflammatory TNFα/NFκB signaling through TRADD, resulting in activation of downstream p-NFκBp65 and cytokine secretion. However, IKKαβ was activated by signals from both the TRADD and PI3K pathways to activate NFκBp65. The loss of CFTR also led to activation of the CFTR-inhibited TRPC6 calcium channel, which may have been further activated by diacylglycerol (DAG), generated by phospholipase C-3β, using the PIP2 substrate provided by PI3K. The elevation in cytosolic calcium due to activated TRPC6 led to an uncoupled state in mitochondria, thus causing an elevation of ROS and resultant oxidative stress. The TRPC6 calcium channel blocker BI 749327 was able to prevent production of ROS and genesis of oxidative stress by denying entry of Ca^2+^. Unexpectedly, BI 749327 also prevented loss of CFTR, thus also suppressing TRADD signaling to NFκB. Finally, essential features of this mechanism, including reduction in β2AR and CFTR and elevation of IL6 and TRPC6 expression, was replicated by exposure of ex-vivo cultures from ferret lung to either PM2.5 or B[a]P. Based on these results from parallel experiments with PM2.5 and B[a]P, we have concluded that the PM2.5 toxin benzo[a]pyrene induces inflammation and oxidative stress in the airway by up-regulation of TRPC6 and inactivation of β2AR/CFTR signaling. *These discoveries mark the first identification of the mechanism by which PM2.5, through its adsorbed toxin B[a]P, can induce inflammation and oxidative stress in lung epithelia*.

**Figure 10:**
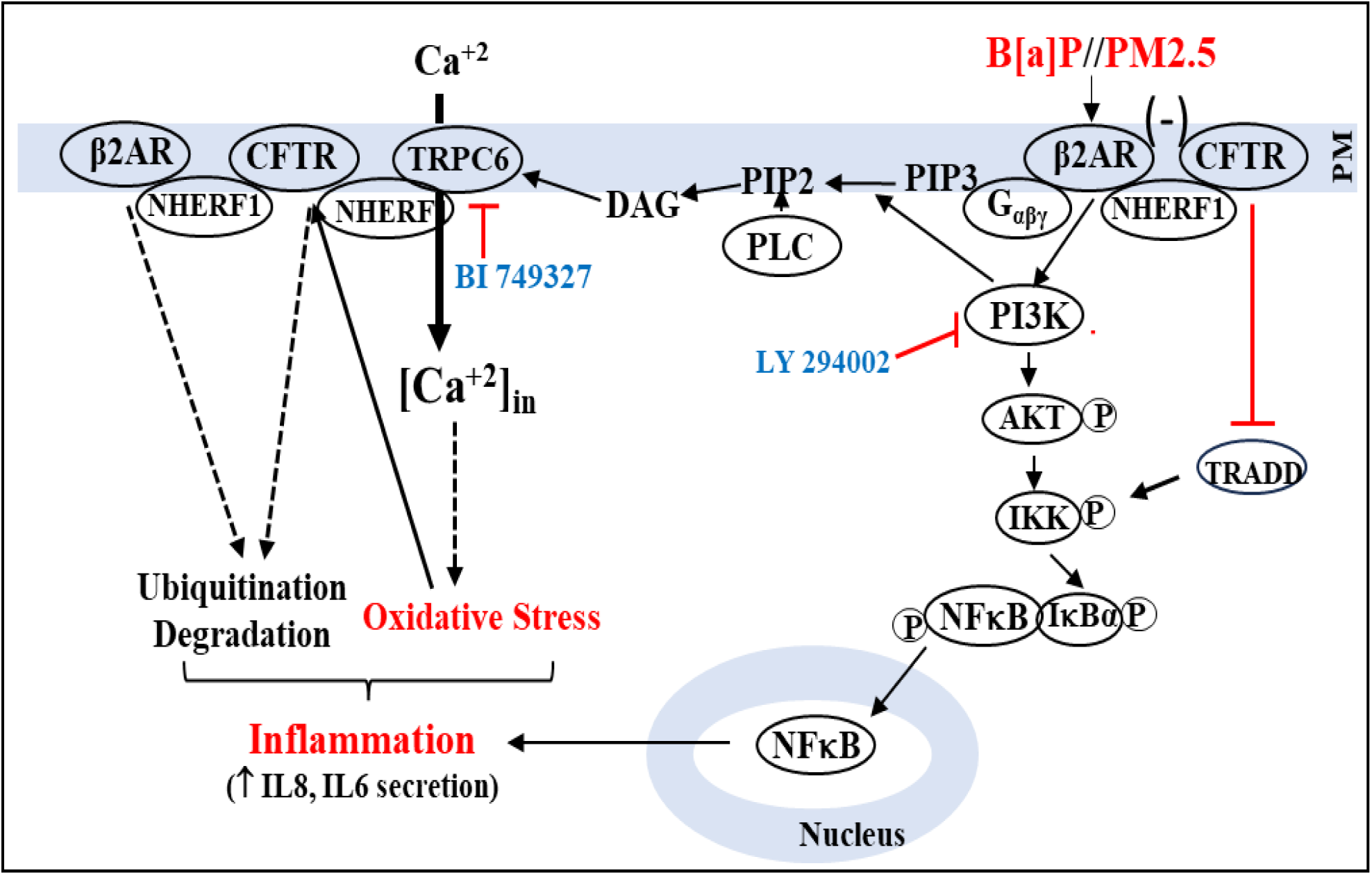
PM 2.5 or the PM 2.5 toxin B[a]P initially acts as an irreversible β2AR agonist to hyperactivate the PI3K signaling pathway. PLC, acting on PIP2, releases DAG. DAG activates the TRPC6 channel to bring extracellular Ca^+2^ into the cell. Increased Ca^+2^ is taken up by mitochondria, which release Reactive Oxygen Species (ROS) and contribute to oxidative stress. BI 749327 can block TRPC6 channel activity and suppress oxidative stress. Meanwhile, hyper-activation of β2AR leads to endosomal β2AR loss, followed by endosomal loss of CFTR (-). Loss of CFTR results in (i) activation of TRADD and downstream activation of IKKαβ to drive NFκB; and (ii) further activation of CFTR-bound TRPC6. NFκB enters the nucleus and drives expression of proinflammatory cytokines IL8 and IL6. Both TRADD and PI3K pathways may be required for optimal NFκB activation, because LY294002 can block inflammation.

### β2AR-CFTR signalplexes and biological condensates

Conventionally, CFTR experiments are designed based on simple linear signalplex models such as [β2AR-NHERF1-CFTR_2_-NHERF1-TRPC6]. However, new understanding suggests that the simple CFTR signalplex structures may occur embedded in multiple large scale clusters on the apical plasma membrane that are much more complex than previously thought [88]. The large-scale CFTR clusters are termed biological coacervates or condensates. Evidence supporting the condensate model includes the calculation that CFTR occurs in dense, discrete clusters on the epithelial plasma membrane that are much larger than might be expected from random diffusion. Further evidence is that biochemical reconstitution of purified CFTR and NHERF1 form similar structures *in vitro.* These structures were found to increase in abundance with phosphorylation and high cholesterol, and to rapidly dissolve upon removal of cholesterol or scaffold proteins such as NHERF1. It has been proposed that the function of cholesterol may be to promote the formation, size and stability of the condensate by favoring CFTR partitioning into cholesterol-rich ordered lipid domains in the apical plasma membrane [89]. The function of NHERF1 includes anchoring the CFTR signalplex to cytoskeletal actin through RhoA and ezrin [90]. The function of ezrin includes increasing the binding abilities of the PDZ domains in NHERF1 [91]. The C-terminal PDZ domain in CFTR, which interacts with NHERF1, may also function to ensuring apical membrane localization of CFTR [91]. In addition, older data [92] and more recent data [88] have converged to indicate that multiple CFTRs may represent hubs in a localized and dynamic association of multiple membrane proteins, many of which may interact with CFTR and with each other through the PDZ linkages provided by scaffold proteins. The scaffold proteins include (i) NHERF1 and (ii) NHERF2/E3KARP (<u>two</u> conserved PDZ domains each) [93–95]; (iii) PDZK1/CASP70 (<u>four</u> conserved PDZ domains [96]); and (iv) Zonula Occludens-1/ ZO-1 (<u>three</u> conserved PDZ domains [97]). In addition, PDZK-1 and ZO-1 can simultaneously link to each other as well as to multiple other PDZ-domain-containing proteins in the condensate, including β2AR [60, 98], ACE2 [75, 99], TRPC6 [62, 100], CAL [101], ezrin [90], NHERF1/EBP50 [102], NHERF2/E3KARP [101], Shank2 [103], PDZK1/CASP70/ NHRF3 [104], ZO-1 [105], and PLC-β3 [106]. Interestingly, PLC-β3, which plays a role in providing DAG from PIP3 to help activate TRPC6 is bound within the CFTR condensate to the first conserved PDZ domain on PDZ1 [106]. *The existence of CFTR and its binding partners in condensates, coavervates or clusters of multiple interacting proteins through PDZ linkages thus complicates simple interpretation of interaction data*.

Finally, there is a dynamic character to the condensate that adds further complexity to understanding the biology of CFTR. Many of these proteins, specifically including CFTR, β2AR and TRPC6, are continuously being endosomally recycled [76]. The purposes of the recycling process include removal of damaged proteins and the return of intact proteins to the membrane. Their rates of endosomal recycling vary. For example, under resting conditions, CFTR recycles through endosomes every 20 minutes, while β2AR recycles every 5 minutes [76]. ACE2 undergoes recycling in 1-4 hours in airway epithelial cells [107]. TRPC6 recycles, but the kinetics have not been determined [108]. However, once a protein within the condensate is activated, its recycling activity is temporarily paused. Thus, within lung epithelial apical membranes there are multiple CFTR-containing condensates in which β2AR, CFTR, TRPC6 and ACE2 are independently recycling in and out of the membrane at different rates, and then reassembling, perhaps in the same or different condensate. Since the TRPC6 inhibitor BI 749327 can rescue CFTR from endosomal loss, it therefore cannot be excluded that blocking the TRPC6 active site puts both TRPC6 and CFTR in a “paused” state within the CFTR condensate(s). *From these considerations it is apparent that the space and time-dependent complexities supporting function in the CFTR condensates represent an important gap in our knowledge of how the dynamics of β2AR, CFTR, and TRPC6 in lung epithelial cells, or elsewhere, are regulated*, *and therefore how PM2.5 and B[a]P might interact with the system*.

### PM2.5, B[a]P, microRNAs and epigenetics

Mechanistically, the ability of PM2.5 and its toxin B]a]P to affect microRNA expression suggests that these agents may also affect gene expression indirectly through epigenetic controls. Mechanistically, five histone proteins intimately associate with nearly the entire genome by forming conditionally stable nucleosome complexes with consecutive links of 145-147 bp of DNA wrapped in 1.7 turns of total nuclear DNA. This nucleosome complex of histones and DNA thereby enables modulation of both chromatin architecture and regulatory accessibility to transcription factors across most genes [109]. Each nucleosome is formed by an octamer of four core histones, H2A, H2B, H3 and H4, and a linker histone H1 or variant [110]. However, when linker H1 is included in a chromosome, the average protected DNA link may be about 168 bp Epigenetic control is then manifest by enzymes that add or subtract covalent chemical marks to DNA, to histones or to other proteins that either activate or suppress gene expression without changing the DNA sequence [111]. MicroRNAs are integral contributors to the epigenetic control process, and are, in turn, also regulated by other epigenetic mechanisms [112]. In addition, because each microRNA can control many transcripts, the possibilities are profoundly increased for regulation of all components of the epigenetic network [113, 114] (see **Fig 11**). The direct relevance of epigenetic mechanisms for PM2.5/B[a]P action on gene expression is thus specifically emphasized by the concordant reduction of the same five B[a]P-dependent microRNAs, *in vitro,* (see **Fig8**), and, *in vivo,* [81] in sera from service members exposed to PM2.5 from outdoor burn pits Consistently, metabolomic evidence, as previously mentioned, showed that the service members who were exposed to PM2.5 also inhaled B[a]P because both B[a]P and the AhR-mediated oxidation product BDPE were found in their sera [84, 85]. A specific example of epigenetic contribution by PM2/5/B[a]P is the case with miR-26a-5p, which targets TRPC6 mRNA. Thus, loss of miR-26a-5p, due to either PM2.5 or B[a]P, can cause elevation of TRPC6 expression [115]. Activated TRPC6, now uninhibited due to endosomally lost CFTR, is made available to mediate influx of calcium, elevate ROS and induce oxidative stress. An additional epigenetic consequence of PM2.5/B[a]P-dependent reduction of miR-26a-3p is that the epigenetic regulator EZH2 is also elevated [116, 117]. EZH2 is one of two catalytic subunits of PRC2 (Polycomb repressive complex 2), that acts to increase the density of H3K27me3 at promoter/enhancer sites on target genes. Importantly, the histone H3 is almost always poised near all possible genes because of the fundamental structure of the nucleosome (*see above*). Thus, increased EZH2 expression, mediated by B[a]P-induced loss of miR-26a-5p, can act promptly to enhance transcription suppression [118]. Separately, B[a]P-reduced expression of miR-766a-3p may also increase expression of EZH1, one of the top 50 Myannnlab targets for miR-766a-3p [119]. Finally, B[a]P-dependent reduction of miR-30b-5p may also be driven by inducing a hypermethylated state of the parent MIR30B locus [120, 121]. This is also caused by increased levels of EZH2 and by elevated levels of methylated CpG on newly synthesized strand of DNA by DNMT1 (maintenance methyltransferase), and by toxin-dependent epigenetic effects that can be stably transmitted across generations by DNMT3B/ 3A (de novo methyltransferases). Thus, increased levels of EZH2 and EZH1 from miR-26a-3p and miR-766a-3p, respectively, may contribute epigenetically to reduction of miR-30b-5p. As further emphasized by **Figure 11**, multiple microRNAs included in the network, including miR-101, miR-26a/b, miR-214, miR-138, and miR-124, are well-documented to directly target EZH2 thereby positioning EZH2 as a central hub for PM2.5/B[a]P dependent post-transcriptional dysregulation. By contrast, EZH1 is largely absent from established miRNA regulatory circuits, consistent with its reduced representation in the network. *This selective enrichment of miRNA-EZH2 interactions suggests that PM2.5 and benzo[a]pyrene (BaP) may preferentially modulate the EZH2-PRC2 axis, leading to enhanced H3K27 trimethylation and epigenetic transcriptional repression of target genes involved in inflammation and immune regulation*.

**Figure 11.**
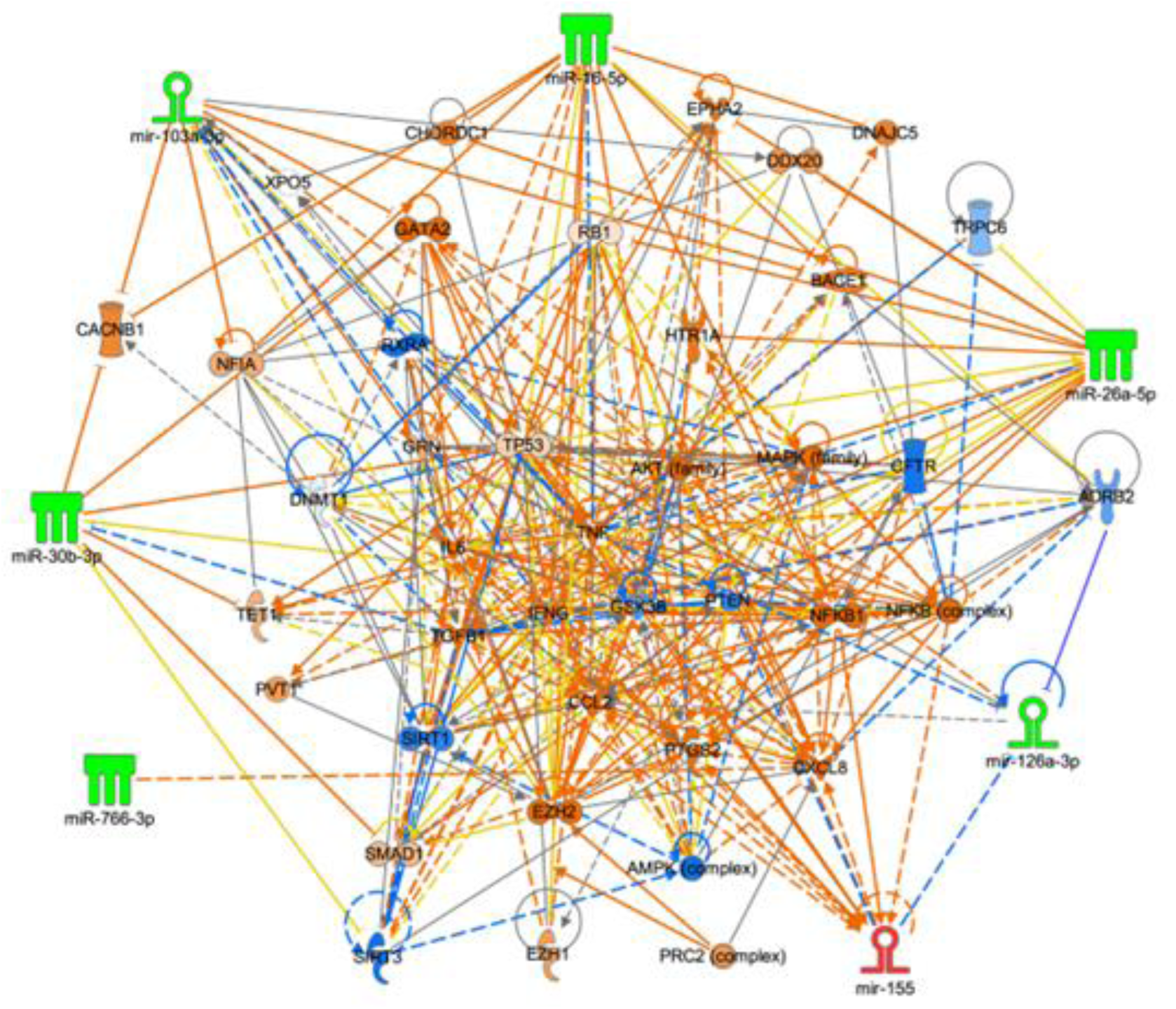
Relationship among microRNAs and targets that are modified by exposure to PM2.5 and B[a]P. Networks were generated using Ingenuity Pathway Analysis, incorporating microRNAs identified as significantly dysregulated following B[a]P exposure *in vitro* and in sera from individuals exposed to PM2.5. The connections depict potential regulatory relationships between these microRNAs and their experimentally validated or predicted targets and associated signaling pathways. The color scheme reflects the direction and magnitude of differential expression. Node shapes correspond to the type and functional category of each molecule (see **Supplemental Figure S5)**. Solid lines indicate direct interactions, whereas dashed lines represent indirect interactions.

In addition to microRNAs, B[a]P] dependent reduction of miR-30b-5p also results in elevation of another kind of epigenetic modifier, long-non-coding RNA PVT1 (lnc RNA Plasmacytoma Variant Translocation 1). PVT1 depletion has been shown (i) to reverse inhibition of miR-30b-5p; (ii) to restore viability; (iii) to reduce apoptosis; and (iv) to suppress proinflammatory cytokine production [122]. Yet another epigenetic example is the manner by which loss of miR126a-3p may occur following activation of β2AR signaling. Mechanistically, activated β2AR, which occurs when B[a[P binds to β2AR [59], activates the ERK1/2 pathway through the G_i_ protein to inhibit AMPKα/p53 signaling. Inhibition of AMPKα/p53 signaling prevents posttranslational maturation of the miR-precursor pre-miR-126 to mature miR-126a-3p [123]. Thus, this mechanism of epigenetic control is by regulating conversion of a microRNA from a precursor transcript to a mature functional state. Finally, B[a]P-dependent reduction of miR-103a-3p has been traced to direct effects of ROS in alveolar epithelial cells that have been generated in people by smoking cigarettes, or in lung epithelial cells treated with cigarette-smoke-extract (CSE), or by directly treating cells with H_2_O_2_, a model ROS [124]. Cigarette smoke extract also activated proinflammatory, NFκB-driven cytokine release [125]. Not surprisingly, cigarette smoke is a source of not only B[a]P, but also the B[a][P carrier PM2.5 [126], and potently reduces CFTR expression [127]. *We conclude from this analysis that epigenetic mechanisms may also contribute to lung pathology in people exposed to PM2.5 and therefore to the PM2.5 toxin B[a]P*.

### Candidate therapeutics for PM2.5/B[a]P toxicity

The PM2.5/B[a]P mechanisms derived so far from studies of lung epithelial cell organoids may be instructive for developing candidate therapeutics. Possible approaches may be found by considering (i) inhibitors of PI3K such as LY1294002; (ii) TRPC6 inhibitors such as BI 749327; (iii) the TRADD-inhibitor digitoxin [128]; and (iv) EZH2 inhibitors. For example, it is possible that further development of PI3K inhibitors such as LY 294002 might lead to development of compounds compatible with human use. In the case of TRPC6 inhibitors such as BI 749327, a close analogue, BI 764198, has been tested in a Phase 1 trial to treat people with Acute Respiratory Distress Syndrome (ARDS) in the ICU [129]. However, deaths within this population compromised determination of safety. The attractive feature of BI 749327 and equivalent analogues is that they have polypharmacy properties. For example, blocking TRPC6 not only inhibited calcium influx, but also rescued CFTR from loss by endocytic recycling. However, the mechanism(s) responsible for these dual drug actions are not fully understood. For example, it is possible that when BI 749327 binds to TRPC6, the inhibitory interaction between TRPC6 and CFTR may be enhanced, and the physiological recycling process may thereby be physiologically paused. It is also possible that NFκB activation may be blocked, not only by the preserved inhibitory action CFTR, but also by reduction in cytosolic calcium increases being blocked by both TRPC6 inhibition, and by consequent reduction in ROS. By contrast, the mechanism for the anti-inflammatory drug digitoxin is much better known. Digitoxin mimics CFTR by inhibiting TRADD, and therefore suppresses the TNFα-NFκB pathway. Digitoxin has been shown in a recent Phase 2a randomized controlled trial (RTC) to be safe for people with CF, and to significantly reduce proinflammatory signaling in upper airway biopsies [130]. Finally, it is possible that clinical grade inhibitors of EZH2 may positively affect the epigenetic network inhibited by B[a]P (see **Figure 11**) [131]. Importantly, at least one EZH2 inhibitor has been approved by the FDA (Tazemetostat/EPZ-66388) and four others are in clinical development [132, 133]). *Consequently, understanding how these drugs may affect components of the pathological PM2.5/B[a]P-driven signaling cascades in the lung may yield insights that could enable future discoveries of candidate therapies for PM2.5/B[a]P exposure*.

### Limitations

This paper has limitations. *First*, it is a limitation that there are other atmospheric pollutants that could be causing pulmonary diseases besides B[a]P that could contribute to enhanced proinflammatory signaling in the lung [134]. However, since PM2.5 and pure B[a]P closely phenocopy each other by causing reduction *in vitro* of all five of the five reduced microRNAs in sera from service members exposed *in vivo* to burn pits, it is likely that B[a]P may be making a significant contribution. *Second*, it is a limitation that the differentiated epithelial organoid used as a target for B[a]P is composed of multiple cell types. However, the same PM2.5 and B[a]P effects on CFTR chloride conductance and protein expression, and on β2AR signaling, were also be detected in the polarized 16HBE14o-cells. The latter cells *do* differentiate to form a tight electrical layer at the air-liquid-interface but *do not* further differentiate into different cell types. *Third*, it is a limitation that we have not experimentally tested the proposed candidate mechanisms of action of the 5 different microRNAs that were reduced by PM2.5 and B[a]P exposure. However, **Figure 11** illustrates predicted interactions, and we suggest that more detailed studies exceed what can be addressed by the present studies, and must be deferred to the future. *Fourth*, it is a limitation that we have not tested β2AR and AhR inhibitors side by side in all the different tests. These proteins clearly have different properties, but may also add to the effects since they are in the same cells and bind B[a]P. We suggest that this complication puts the analysis beyond what can be accomplished in the present studies and must also be deferred to the future. *Fifth*, it is a limitation that we do not know how the TRPC6 inhibitor BI 749327 not only blocks the TRPC6 channel but also preserves its inhibitory signalplex partner CFTR. We suggest that this complication also puts the analysis beyond what can be accomplished in the present studies and must therefore be deferred to the future. *Sixth*, it is a limitation that we do not understand the mechanism by which PI3K and the TRADD-dependent pathways differentially drive NFκB through IKKαβ. However, the PM2.5/B[a]P driven PI3K pathway may be dominant in the presence of either PM2.5 or B[a]P. We suggest that the investigation of this interesting mechanism also puts the analysis beyond what can be accomplished in the present studies and must also be deferred to the future.

### Conclusion

The PM2.5 toxin Benzo[a]Pyrene (B[a]P) is toxic to the lung by virtue of its ability to form an irreversibly activating complex with β2AR, and thus indirectly with CFTR and TRPC-6. The B[a]P:β2AR complex activates Gi, which drives the proinflammatory [PI3K/AKT/IKKαβ/NFκB] signaling pathway. This pathway is blocked by the PI3K inhibitor LY 294002. Desensitization of B[a]P:β2AR occurs by endosomal recycling, which is accompanied by equivalent loss of CFTR in the β2AR-NHERF1-CFTR signalplex. With loss of CFTR, TRADD is activated to drive downstream activation of NFκB signaling, and the CFTR-inhibited TRPC6 calcium channel is itself released, activated and further expressed. Increased cytosolic calcium from TRPC6 leads to mitochondrial dysfunction, increased ROS production and Oxidative Stress, which is blocked by the TRPC6 inhibitor BI 749327. Blocking TRPC6 with BI 749327 also pauses the B[a]P-dependent loss of CFTR and downstream proinflammatory consequences. Thus, the contributions of both activated PI3K and loss of CFTR to proinflammatory NFκB signaling, and increased cytosolic calcium concentration due to activation of TRPC6, results in a proinflammatory cystic fibrosis-like phenotype in persons who have inhaled PM2.5, and thus also have inhaled the adsorbed B[a]P toxin. Studies of *ex-vivo* ferret tracheo-bronchial slice cultures support this conclusion. These discoveries mark the first identification of a mechanism by which exposure to PM2.5, or to the PM2.5 toxin B[a]P itself, can induce inflammation and TRPC6-dependent oxidative stress in lung epithelia.

## Methods

### Materials and Reagents

All chemicals and reagents were purchased commercially. Benzo[a]pyrene, PM 2.5 (ERMCZ110), CFTR_inh_-172, IBMX, amiloride, Forskolin, MG-132, L-glutathione, and Millipore Immobilon Western (Sigma-Aldrich); BI 749327, LY 294002, and LY 303511 (Cayman Chemical); MTT (3-(4,5-Dimethylthiazol-2-yl)-2,5-Diphenyltetrazolium Bromide; EZ-Link™ Sulfo-NHS-SS-Biotin, CellROX green reagent, Dynabeads Protein G, Pierce Streptavidin magnetic beads, Halt™ protease and Phosphatase Inhibitor Cocktail (Thermo Fisher) ; Fluo-4 AM and Probenecid (Invitrogen); Human IL-8/CXCL8 DuoSet® ELISA kit, Human IL-6 ELISA DuoSet® kit, and Human TNF-alpha DuoSet® ELISA (R&D Systems); Ferret IL-6 ELISA kit (Genorise, PA, USA); Pneumacult-ALI media kit and Pneumacult-EX Plus Basal media kit (Stemcell); and Gibco MEM (Thermo Fisher).

All antibodies were purchased commercially. β-actin (Sigma, A5441); CFTR (R&D Systems, MAB1660); CFTR (SCBT, sc-376683); β2AR (Novus Biologicals, NBP261710); TRADD (SCBT, sc-46653); TNFR1 (Cell Signaling, 3736), IKKβ (SCBT, sc-8014); IκBα (Cell Signaling, 9242); phospho-IκBα Ser32/36 (Cell Signaling, 9246); phospho-NFκB p65 Ser276 (Cell Signaling, 3037); NFκB p65 (Cell Signaling, 8242); phospho-IKKα/β Ser176/180 (Cell Signaling, 2697); IKKα (Cell Signaling, 11930); AKT (Cell Signaling, 9272); phosphor-AKT Ser473 (Cell Signaling, 9271); TRPC6 (Protein Tech, 18236-1-AP); and Ubiquitin (Protein Tech, 80992-1-RR).

### Cell Cultures and Treatments

We thank Dr. R.G. Crystal (Cornell Medical College, New York City, NY) for the gift of the hTERT-transformed BCi-NS1.1 basal stem cell. The basal cells were cultured, maintained, and differentiated on Transwell or Snapwell inserts under Air-Liquid Interface (ALI) conditions according to the instructions from Dr. Crystal’s lab. Human bronchial 16HBE14o-(SCC150) cell lines were purchased from Sigma-Aldrich, and were cultured and subdivided according to the manufacture’s instruction. Human BCi-NS1.1 basal stem cell and 16HBE14o-cells were seeded at a concentration of 5×10^5^ cells/cm^2^ on 12-mm Snapwell or Transwell inserts and 9×10^5^ cells/cm^2^ on 24-mm Transwell inserts. All experiments were performed in differentiated BCi-NS1.1 (dBCi) and polarized 16HBE14o-cells after maintaining at the air-liquid-interface (ALI) for 25-28 days and for 9-10 days, respectively, in a humidified tissue culture incubator at 37°C, 5% CO_2_.

The following compounds were dissolved in DMSO: B[a]P, LY 294002, LY 303511, BI 749327, and PM 2.5 powder. For cell treatments, both apical and basolateral sides of cell monolayers were exposed to DMSO (Control), B[a]P, or PM 2.5 in the presence or absence of LY 294002, LY 303511, or BI 749327; after 6-hr incubation, treated media from the apical sides were removed and cells were continued to expose to drugs for additional 18 hrs under ALI conditions. The DMSO concentration in all treatments was less than 1.0%. After treatments, cell monolayers were prepared for cell viability, Ussing chamber, cell surface biotinylation, endosomal recycling, immunoprecipitation, or microRNA assays. The treated media from the basolateral sides were collected for measuring the secreted levels of IL-8, IL-6 and TNFα.

### Cell viability assay

Cell viability was assessed using the MTT assay. To each treatment well of a 12-well plate, an equal volume of MTT solution (5 mg/mL) to the existing media in the culture, and cells were incubated in CO_2_ incubator in the dark for 2 hrs. The medium was removed; Formazan crystals formed by the cells were dissolved using 200 µl of solubilization solution; and the plate was shaken on a rotary to ensure complete dissolution of the crystals. Absorbance was then read at 570 nm using the CLARIOstar microplate reader (BMG LABTECH, UK).

Measurement of IL-8, IL-6 and TNFα

Culture media from the basolateral compartments were collected and used to assay for IL-8, IL6 and TNFα using respective DuoSet® ELISA kits, and performed according to the manufacturer’s instructions. Concentrations of human TNFα, IL-6, and IL-8 were calculated using a standard curve and normalized to the MTT or total protein concentration values.

### Ussing chamber analysis

Treated Snapwell inserts were mounted in Ussing Chambers (Physiologic Instruments, Reno, Nevada). A basolateral-to-apical chloride gradient was imposed by replacing NaCl with sodium Gluconate. The chloride-containing solution was 120 mM NaCl, 20 mM NaHCO_3_, 5 mM KHCO_3_, 1.2 mM NaH_2_PO_4_, 1.2 mM CaCl_2_, 1.2 mM MgCl_2_, and 5.6 mM glucose. In the chloride-free solution, all Cl salts were exchanged for gluconate salts, and Ca^+2^ was increased to 5 mM to compensate for the chelation of Ca^+2^ by gluconate. Both chamber compartments were gassed with 5% CO_2_ (pH 7.4). Experiments were done at 37°C. CFTR channel activity was detected by first inactivating sodium currents with 100 µM amiloride; then activating CFTR chloride channels with 10 µM forskolin and 100 µM IBMX; then specifically inactivating CFTR channels with 10 µM CFTR_inh_-172. At the end of the assays, cells were washed with PBS and lysed in RIPA buffer supplemented with the anti-protease/phosphatase cocktail or labeled with cell impermeable Sulfo-NHS-SS-biotin for Western blot analyses.

### Cell surface biotinylation

After B[a]P treatment, ALI differentiated cells were washed three times with Ca^+2^/Mg^+2^-PBS buffer (pH 8.2) and then incubated with cell impermeable Sulfo-NHS-SS-biotin (1 mg/mL) for 2 hours at 4^0^C in the dark, followed by washing three times with 100 mM Tris buffer to quench all free biotin reagent. Biotinylated cells were lysed in RIPA buffer supplemented with anti-protease/ phosphatase cocktail; biotinylated proteins at 1 mg/mL in the lysates were extracted using streptavidin magnetic beads; and biotinylated β2AR and CFTR were determined by Western blot.

### Endosomal recycling

To measure recycling of endogenous β2AR and CFTR, ALI differentiated cells were first biotinylated 1 mg/ml Sulfo-NHS-SS-biotin for 2 hours at 4^0^C in the dark, followed by incubation at 37^0^C for 6 hours with culture media containing DMSO control or different concentrations of B[a]P. After 37^0^C incubation, biotin molecules remaining at the cell surface were stripped five times with glutathione (GSH) solution (Bomberger JM et al, 2011). One biotinylated cell sample without 37^0^C incubation and GSH stripping and other biotinylated cell sample without 37^0^C incubation and with GSH stripping were used as control for biotin labeling efficiency and control for GSH stripping efficiency, respectively. Biotinylated proteins were extracted from the lysates using streptavidin magnetic beads, and biotinylated β2AR and CFTR were determined by Western blot. β-actin was used in the Western blot for equal amounts of biotinylated samples used in the biotinylated protein extraction.

### CFTR Ubiquitination

Both apical and basolateral sides of polarized 16HBE14o-cell monolayers were exposed to DMSO (Control) or different concentrations of B[a]P in the presence of 20 µM proteasome inhibitor MG-132; then treated media only from the apical side were removed after 6-hr incubation, and cells were continuously exposed to B[a]P further for 18 hrs at the ALI. After treatment, cell monolayers were lysed in RIPA buffer. The lysates (1mg/mL) were precleared with Protein G Dynabeads and then incubated for 6 hrs at 4°C with 2 µg of normal mouse serum or anti-CFTR R Domain antibody, followed by incubation for overnight with Protein G Dynabeads. After washing, CFTR ubiquitination in the immunoprecipitated complexes was analyzed by Western blot with an anti-ubiquitin antibody.

### Fluorescence measurement of intracellular free calcium ([Ca2+]i)

16HBE14o-cells at a density of 1×10^5^ cells/mL grown in black visiplateTM 24-well tissue culture plates were treated with DMSO or different concentrations of B[a]P or PM 2.5 for 24 hrs at 37°C, followed by washing three times with Ca^+2^/Mg^+2^-free HBSS buffer and incubation for 1 hr at 37°C with 10 µM cell permeant Fluo-4 AM and 2.5 mM organic anion-transport inhibitor probenecid diluted in Ca^+2^/Mg^+2^-free HBSS buffer. After washing, the basal fluorescence intensity of Fluo-4 AM in the cells bathed in Ca^+2^/Mg^+2^-free HBSS buffer was recorded at Ex 483 nm/Em 530 nm using the CLARIOstar microplate reader (BMG LABTECH). When the baseline was stable, 1 mM CaCl_2_ was added, and the fluorescent intensity was continued to record over a time period. At least three independent experiments in duplication were conducted.

### Oxidative stress assay

CellROX Green reagent (ThermoFisher) was designed to detect the production of reactive oxygen species (ROS) in living cells. These reagents are cell-permeable and show no or very weak fluorescence in a reduced state, but their oxidation in the presence of ROS results in a strong fluorescence. Both apical and basolateral sides of polarized 16HBE14o-cells were incubated with PM 2.5, B[a]P, or hydrogen peroxide (H_2_O_2_) at 37°C in the CO_2_ incubator. After 6hr incubation, treated media from the apical sides were removed, and replaced with 10 µM CellROX Green in Ca^+2^/Mg^+2^-free HBSS buffer, followed by further incubation for 18 hrs. Cell monolayers were washed three times with PBS at room temperature and lysed in 0.1% Triton X-100-PBS. The cell lysates were transferred to a black, clear bottom 24-well plate, and fluorescence at Ex 483 nm/Em 530 nm was recorded using the CLARIOstar microplate reader (BMG LABTECH). Background fluorescence was subtracted from each value and normalized with the total protein concentration.

### Western blotting

Treated cells were washed with PBS and lysed in RIPA buffer supplemented with the anti-protease/phosphatase cocktail. Equivalent amounts of lysates (50 µg/sample) were electrophoresed on 4-12% gradient gels (Invitrogen), transferred to PVDF membranes, and membranes were probed at 4^0^C overnight with corresponding primary antibody diluted to the concentration according to the manufacture’s recommended instructions. After incubation with appropriate secondary antibody coupling to HRP and washing, immunoreactive bands were exposed to chemiluminescent HRP substrate solution and visualized using BioRad ChemiDocTM Imaging System and quantified using the Imagelab software.

### RNA isolation and measurement of microRNAs

Both apical and basolateral sides of differentiated dBCi cells were incubated with DMSO or different concentrations of B[a]P for 24 hours. Treated media from the apical sides were removed after 6-hr incubation, and cell treatment continued further under ALI conditions. The cells were lysed and total RNA were extracted using mirVana total RNA isolation kits (Life Technologies, Frederick, MD) following the manufacturer’s protocol. RNA yield and quality was analyzed on a NanoDrop spectrophotometer ND-1000 (Thermo Fisher Scientific Inc., Rockville, MD) with all samples showing A260/A280 absorbance ratios between 1.8 and 2.0, indicative of high-quality RNA. MiRNAs analysis was performed with specific Taqman probes and U6 was used as an endogenous control to normalize expression of miRs. The data represents average of three technical replicates. The relative fold changes of miRs were analyzed using the 2^-ΔΔCT^ method. The data represents relative fold changes with standard error of the mean (mean ± SEM).

### Pathway analyses

In silico analysis of differentially expressed miRNAs following B[a]P exposure *in vitro* and in sera from individuals exposed to PM2.5 was conducted using Ingenuity Pathway Analysis (IPA, Qiagen; https://www.qiagenbioinformatics.com/products/ingenuity-pathway-analysis). Differentially expressed miRNAs and their experimentally validated or predicted targets were further analyzed to construct functional interaction networks involving regulatory molecules and signaling pathways. Fisher’s exact test was used to evaluate the enrichment of differentially expressed miRNAs within specific biological pathways. A significance threshold of p < 0.05 was applied unless otherwise stated.

### *Ex vivo* ferret lung culture model

We thank Dr. Sharon Juliano (USUHS, Department of Anatomy, Physiology and Genetics) for donating lung tissues from two ferrets. Lung parenchymal tissues were washed three times with PBS supplemented 1X penicillin/streptomycin in a petri dish, and the tissues were then dissected into small cubes, 9 cubes per ferret lung with the average wet weight of 400 mg, containing tracheobronchial tubes. The tissues were transferred to a 24-well tissue culture plate, conditioned for 1hr in Pneumacult-EX Plus Basal media cultured, and then cultured for 24 hr in the same media containing DMSO, 100 ng/ml PM 2.5, or 10 µM B[a]P in a humidified tissue culture incubator at 37°C, 5% CO_2_. After incubation, culture medium samples were collected, centrifuged at 400 × g, 4°C for 5 min to remove cellular material, and analyzed by Ferret IL-6 ELISA kit. Treated tissues were washed with PBS and homogenized in RIPA buffer supplemented with anti-protease/phosphatase inhibitor cocktail using a handheld motor driven tissue homogenizer, followed by centrifugation at 20,000 rpm for 5 min. Equivalent amounts of lysates (50 µg/sample) were electrophoresed on 4-12% gradient gels and transferred to PVDF membranes, and membranes were probed at 40C overnight with corresponding primary antibody for CFTR (CFF Antibody Distribution program, mAb 660 + mAb 596), β2AR (Novus Biologicals, NBP261710), TRPC6 (Protein Tech, 18236-1-AP), and β-Actin (Cell Signaling, 4970).

### Statistics

Except as noted, all plotted data points are the means of 3 or more independent experiments. Results are expressed as mean ± SD. Statistical differences among treatment groups were analyzed by one-way ANOVA using Tukey as a post-hoc test for multiple comparisons. *p* < 0.05 is considered significant.

## Acknowledgements

We gratefully acknowledge support for this project by NIH Grant 1RO1HL167048 (Pollard HB, PI). We thank Patty Deuster for insightful comments. We thank Laiman Tavedi for expert administrative support. We thank high school summer intern Elsa Clark for help in data curation. USUHS Clearance number: REQ0081346.

## Disclaimer

The opinions, interpretations, conclusions and recommendations are those of the authors and are not necessarily endorsed by the U.S. Army, Department of War, the U.S. Government or the Uniformed Services University of the Health Sciences. The use of trade names does not constitute an official endorsement or approval of the use of such reagents or commercial hardware or software. This document may not be cited for purposes of advertisement.

## Contributions

HC, MO, RB, SJ, NF, TF and HP designed experiments. HC, MO, WR, OL, SD, TC, QY and TD performed experiments. HC, HP, MO, TC, SJ, RB and BP contributed to data analysis. HP, HC, MO, SJ, RB, TF, and BP wrote the paper. All co-authors critically read the paper.

## Data availability

The data that support the findings of this study are available from the corresponding author upon reasonable request.

## SUPPLEMENTAL DATA

**Supplemental Figure S1:**
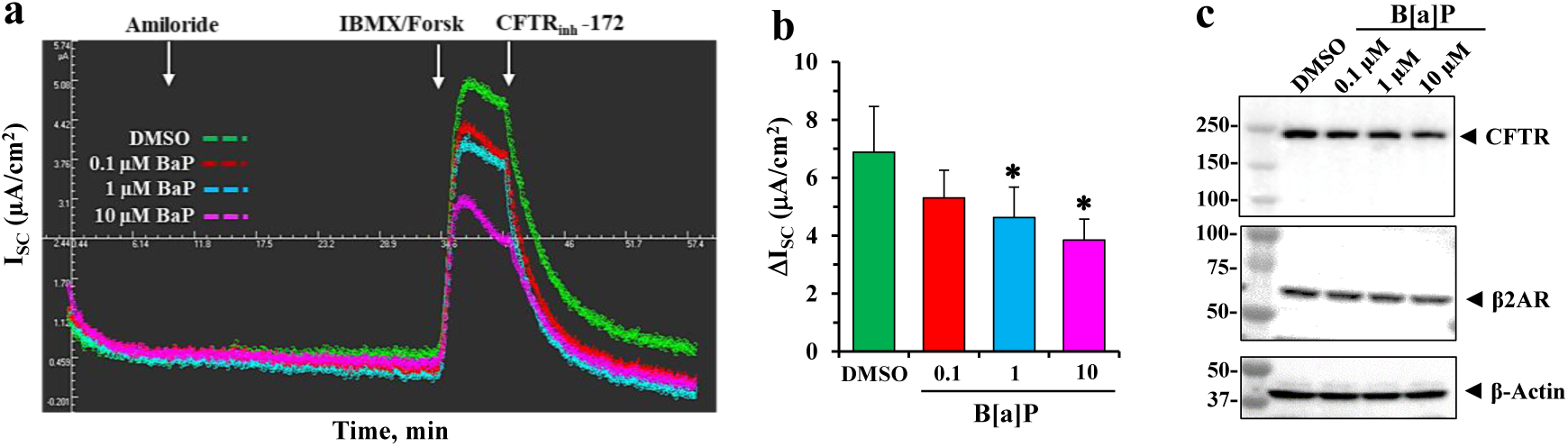
Effects of Benzo[a]Pyrene (B[a]P) on functional expression of CFTR in polarized 16HBE14o cells. Both apical and basolateral sides of polarized 16HBE14o-cells (ALI for 8-10 days) were incubated with different concentrations of B[a]P. Treated media from the apical sides were removed after 6-hr incubation, and cell treatment continued further for 18 hrs under ALI conditions. (**a**) Representative I_sc_ tracings of CFTR-dependent short-circuit currents (I_sc_) measured in Ussing chambers as the changes in response to Amiloride, IBMX/Forskolin, and CFTRinh-172. (b) Summary of changes in I_sc_ is shown. Data represent the mean of three independent experiments, and each experiment was performed in duplication (n = 3). Statistical *p* values were determined with a one-way ANOVA, followed by the Tukey post-hoc test comparing the B[a]P treatments to the DMSO control; * *p* < 0.05; ** *p* < 0.01. (**c**) Representative Western blot images of β2AR and CFTR expression levels in treated cells after Ussing chamber analyses in (**a**).

**Supplemental Figure S2:**
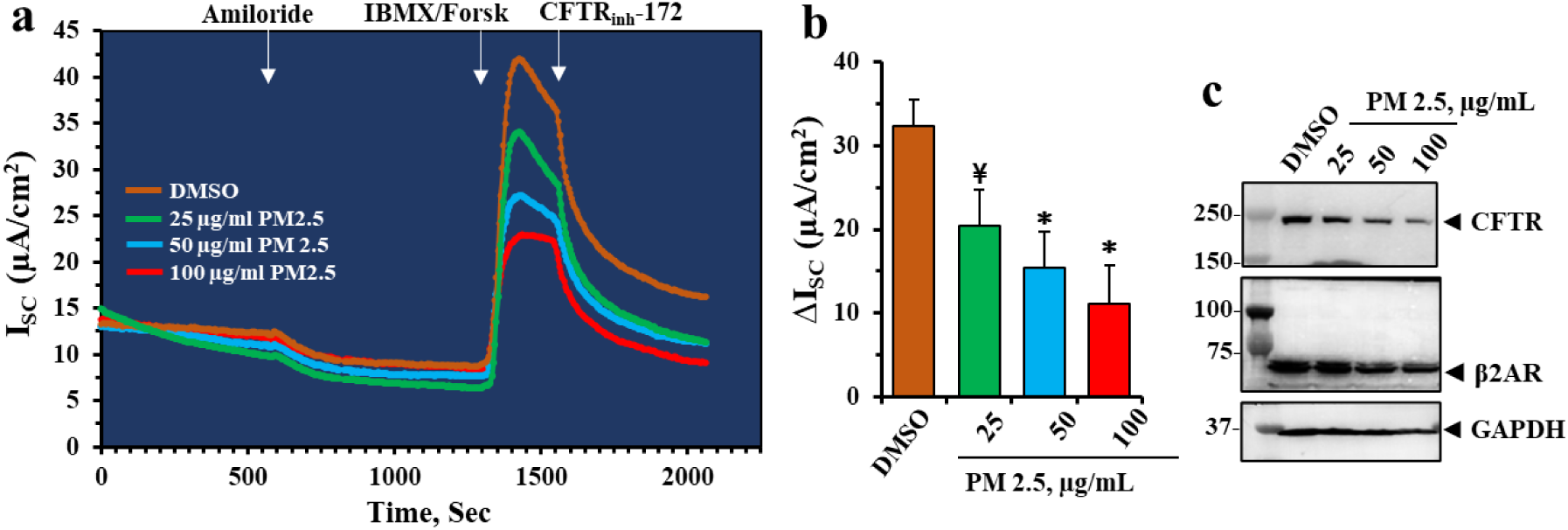
Effects of PM 2.5 on functional expression of CFTR in polarized 16HBE14o cells. Both apical and basolateral sides of polarized 16HBE14o-cells (ALI for 8-10 days) were incubated with different concentrations of PM 2.5. Treated media from the apical sides were removed after 6-hr incubation, and cell treatment continued further for 18 hrs under ALI conditions. (**a**) Representative I_sc_ tracings of CFTR-dependent short-circuit currents (I_sc_) measured in Ussing chambers as the changes in response to Amiloride, IBMX/Forskolin, and CFTRinh-172. (b) Summary of changes in I_sc_ is shown. Data represent the mean of three independent experiments, and each experiment was performed in duplication (n = 6). Statistical *p* values were determined with a one-way ANOVA, followed by the Tukey post-hoc test comparing the B[a]P treatments to the DMSO control; ¥, *p* < 0.05; *, *p* < 0.01. (**c**) Representative Western blot images of β2AR and CFTR expression levels in treated cells after Ussing chamber analyses in (**a).** All Western analyses represent the results of three independent experiments, and β-actin was used for equal loading of protein.

**Supplemental Figure S3.**
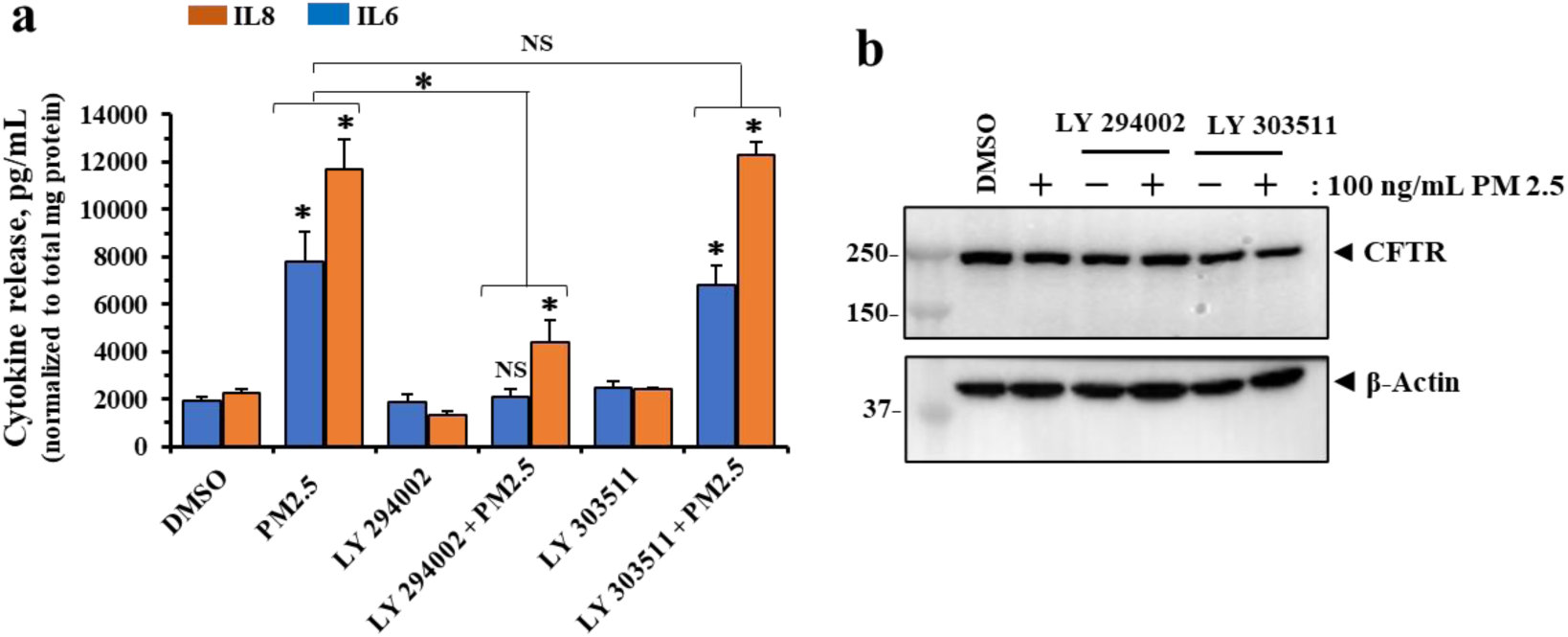
PM 2.5 induces cytokine secretion via the PI3K signaling pathway in polarized 16HBE14o-cells. ALI cells were treated with DMSO, 100 ng/mL PM 2.5, 10 µM LY 294002 ± PM 2.5, or 10 µM LY 303511 ± PM 2.5 for 24 hrs. (**a**) IL-6 and IL-8 from the basolateral sides were determined using the DuoSet® ELISA kits. Data are presented as the mean ± SD (n=3). Statistical *p* values were determined with a one-way ANOVA, followed by the Tukey post-hoc test comparing all treatments to the DMSO control, or PM 2.5 to PM 2.5 ± LY 294002 or LY 303511; *, *p* < 0.01. (**b**) Representative Western blot images showing CFTR and β-actin expression. β-actin was used to ensure equal loading of protein. All data were obtained from three independent experiments, and each experiment was analyzed in duplication.

**Supplemental Figure S4.**
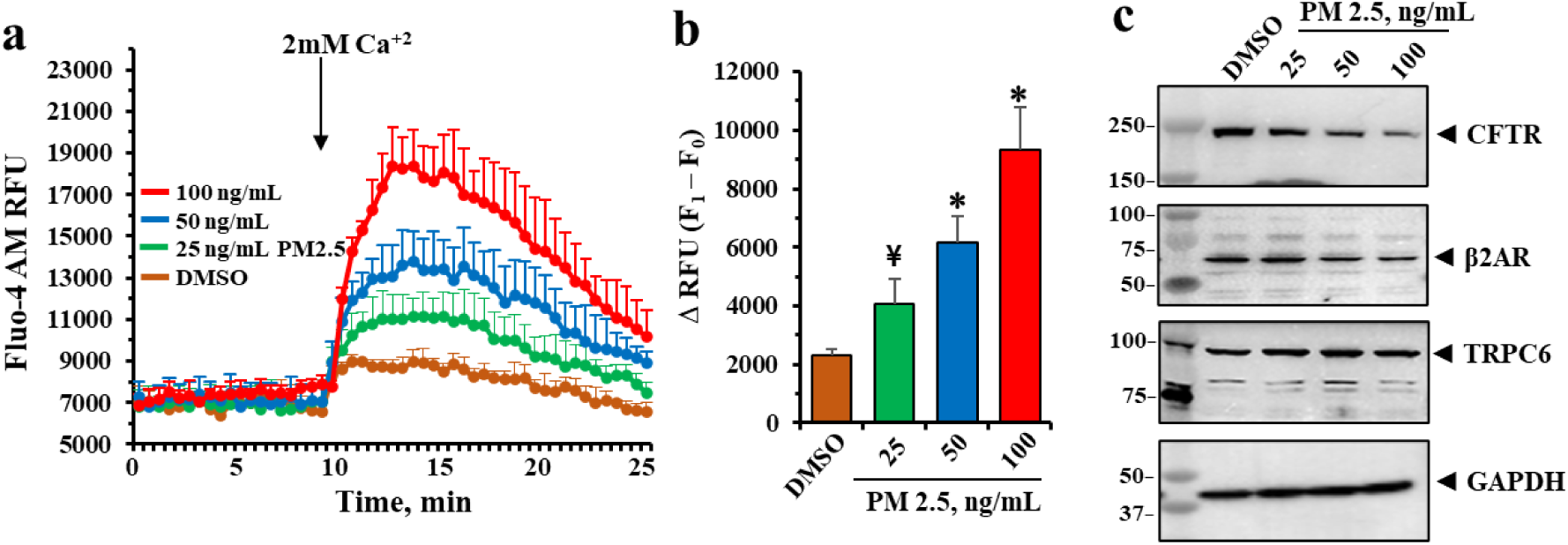
PM 2.5 induces calcium influx in 16HBE14o-cells. (**a**) After 24 hr treatment, fluorescence intensity traces obtained using the fluo-4 AM when DMSO- or PM 2.5-treated 16HBE14o-cellswere stimulated with assay buffer containing 2 mM Ca. Each trace is the mean of three independent experiments with standard deviations (vertical bars). (**b**) Corresponding histograms summarize the mean area under the curve (AUC) after stimulation by 2 mM Ca. (**c**) Representative Western blot images of CFTR, β2AR and TRPC6 expression levels in treated cell in (**a**). All data were obtained from four independent experiments. Statistical *p* values were determined with a one-way ANOVA, followed by the Tukey post-hoc test comparing the PM 2.5 treatments to the DMSO control; ¥, *p* < 0.05; *, *p* < 0.01.

**Supplemental Figure S5.**
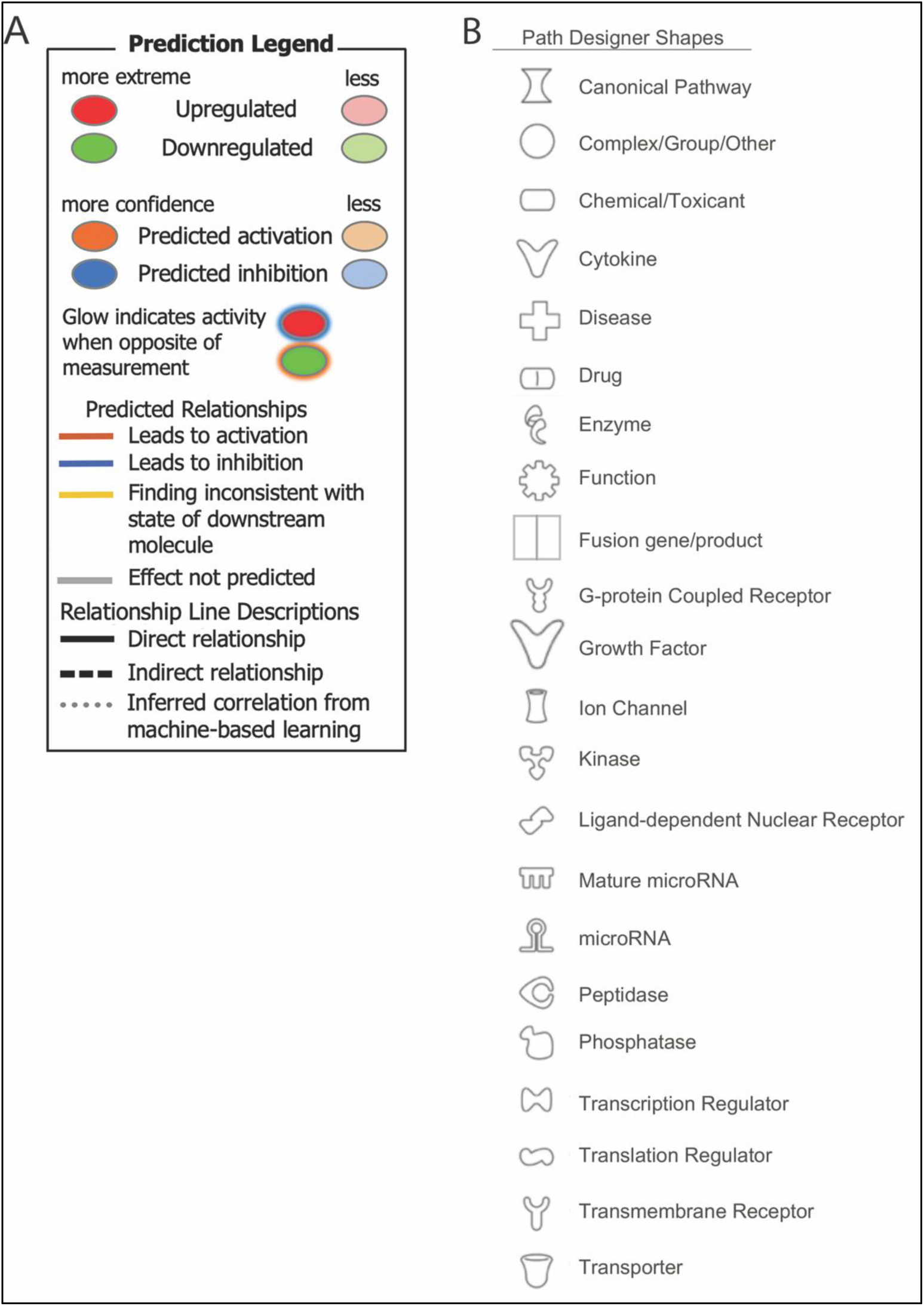
Prediction Legend for Ingenuity Pathway Analysis, Qiagen.com

